# Genomic, transcriptomic and proteomic depiction of iPSC-derived smooth muscle cells as emerging cellular models for arterial diseases

**DOI:** 10.1101/2022.05.01.490058

**Authors:** Lu Liu, Charlène Jouve, Joséphine Henry, Takiy-Eddine Berrandou, Jean-Sébastien Hulot, Adrien Georges, Nabila Bouatia-Naji

**Author notes:** These authors co-supervised this study (Adrien Georges, Nabila Bouatia-Naji).

## Abstract

**Background:** Vascular smooth muscle cells (VSMCs) plasticity is a central mechanism in cardiovascular health and disease. We aimed at providing deep cellular phenotyping, epigenomic and proteomic depiction of SMCs derived from induced pluripotent stem cells (iPSCs) and evaluating their potential as cellular models in the context of complex genetic arterial diseases.

**Methods:** We differentiated 3 human iPSC lines using either RepSox (R-SMCs) or PDGF-BB and TGF-β (TP-SMCs), during the second half of a 24-days-long protocol. In addition to cellular assays, we performed RNA-Seq and assay for transposase accessible chromatin (ATAC)-Seq at 6 time-points of differentiation. The extracellular matrix content (matrisome) generated by iPSCs derived SMCs was analyzed using mass spectrometry.

**Results:** Both iPSCs differentiation protocols generated SMCs with positive expression of SMC markers. TP-SMCs exhibited greater capacity of proliferation, migration and lower calcium release in response to contractile stimuli compared to R-SMCs. RNA-Seq data showed that genes involved in the contractile function of arteries were highly expressed in R-SMCs compared to TP-SMCs or primary SMCs. Matrisome analyses supported an overexpression of proteins involved in wound repair in TP-SMCs and a higher secretion of basal membrane constituents by R-SMCs. Open chromatin regions of R-SMCs and TP-SMCs were significantly enriched for variants associated with coronary artery disease and blood pressure, while only TP-SMCs were enriched for variants associated with peripheral artery disease.

**Conclusions:** Our study portrayed two iPSCs derived SMCs models presenting complementary cellular phenotypes of high relevance to SMC plasticity. In combination with genome-editing tools, our data supports high relevance of the use of these cellular models to the study of complex regulatory mechanisms at genetic risk loci involved in several arterial diseases.

**Graphical Abstract:** 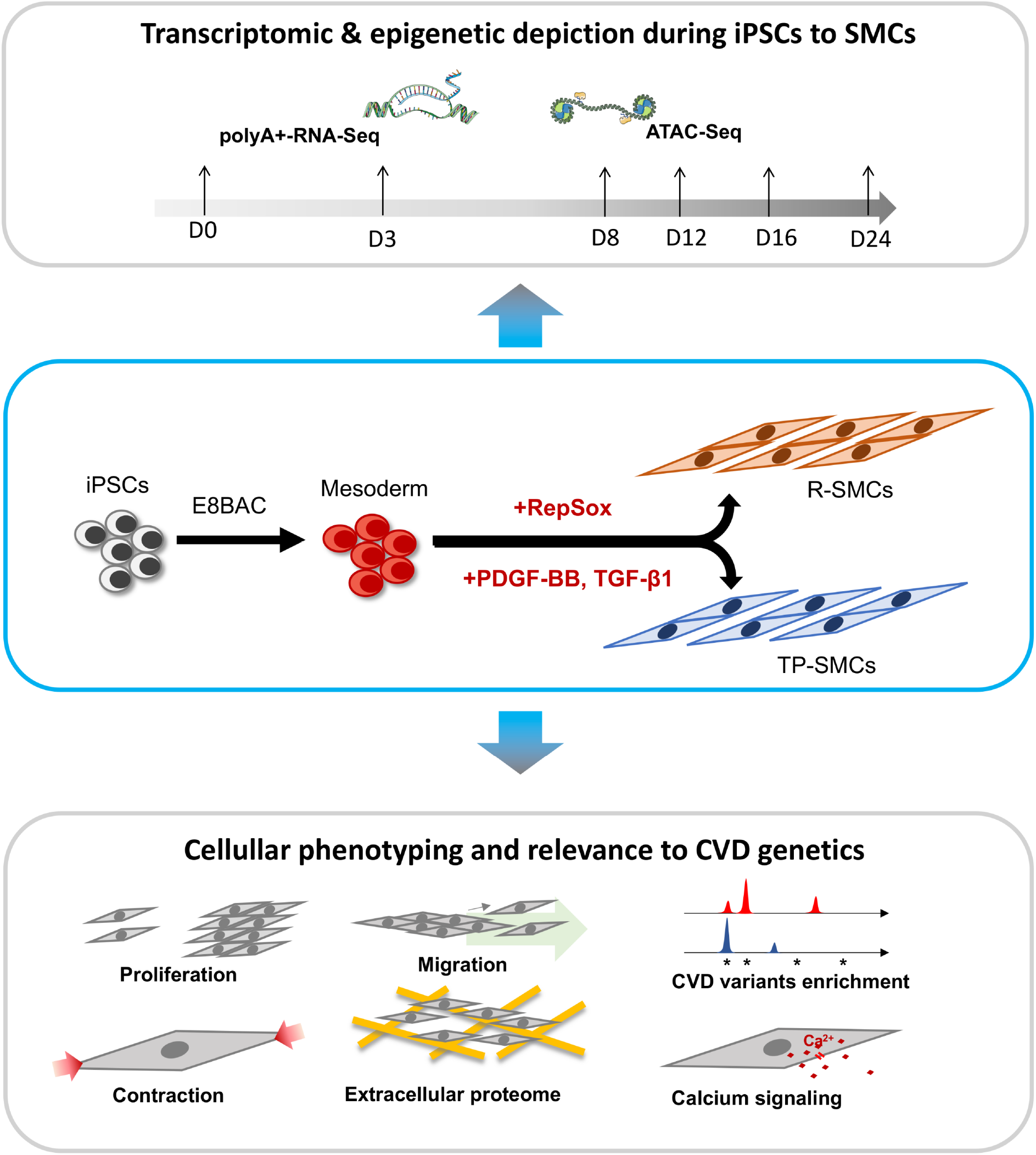

## Introduction

Vascular smooth muscle cells (VSMCs) are contractile differentiated cells located in the mature medial layer of arteries. They are the main structural components of muscular arteries, which distribute blood flow in organ and tissues^1^. The contraction of VSMCs is essential to maintain vascular tone and is mainly mediated by the release of Ca^2+^ into the cytoplasm by influx from the extracellular space and/or release from the sarcoplasmic reticulum^2^. Mature contractile SMCs hold a capacity to switch phenotype to a dedifferentiated or synthetic state characterized by an increasing capacity of proliferation, migration and extracellular matrix remodeling. In response to environmental alterations, synthetic VSMCs play an important role in particular during vascular wound repair and in disease^3^. This process was shown to be regulated by various stimuli such as platelet derived growth factor-BB (PDGF-BB), transforming growth factor beta (TGF-β), oxidized phospholipids and retinoic acid^3-5^.

The modulation of SMC phenotype plays a major role in cardiovascular disease, a leading cause of mortality and morbidity worldwide^6^. In atherosclerosis, VSMCs recruited from the vessel wall contribute to the formation of atherosclerotic plaques and play both protective and detrimental roles on plaque buildup and stability^7, 8^. In hypertension, vascular remodeling capacity is highly dependent on VSMCs and critically regulated by their plasticity to undergo phenotypic switching^9, 10^. VSMCs function is also reported as target of genetic defaults that affect the structural integrity of the vascular wall through impaired extracellular matrix (ECM), not only in rare vascular diseases such as Marfan and vascular Ehlers-Danlos syndromes^11, 12^, but also in the context of common and complex diseases, including coronary artery disease and hypertension^13-15^. In this case, common and non-coding variants are more likely to affect transcriptional regulation of target genes^16^. Risk variants are often located in regulatory elements mobilized in a specific tissue and cell state, which is a challenge for *in vitro* studies. The generation of cellular models mimicking the epigenetic properties of contractile VSMCs in mature vessels is critical to the study of a large proportion of genetic loci implicated in cardiovascular disease. Primary human SMCs present several limitations to model contractile functions given that major alterations of their regulatory profile are reported *in vitro*, in addition to important technical difficulties to modify them genetically^16, 17^. Differentiation of SMCs from genetically editable human induced pluripotent stem cells (iPSCs) in a controlled environment emerged in the past years as a promising approach to obtain genetic and cellular models for human diseases involving arterial wall remodeling ^18, 19^.

Following the development of iPSCs, multiple differentiation protocols have been proposed to generate SMCs and reflect the wide diversity of embryologic origins for vascular SMCs^20-22^. For example, protocols inducing mesodermal differentiation, either lateral plate or paxaxial, are typically used to model coronary artery and descending aorta, respectively, while neural-crest precursors are used to model ascending aorta and cerebro-cervical arteries^20-22^. The majority of these differentiation protocols make use of platelet-derived growth factor-BB (PDGF-BB) and transforming growth factor-β1 (TGF-β1) to induce SMC phenotype^20, 22, 23^. These protocols lead to the production of cells robustly expressing SMC marker genes. However, the resulting iPSCs-derived SMCs generally expressed variable levels of markers specific for contractile function, mainly smooth-muscle myosin heavy chain (smMHC, encoded by *MYH11*)^24-27^. Recently, a small molecule initially identified as an inhibitor of TGF-β signaling (RepSox) was found to lead to increased expression of contractile markers such as smooth-muscle myosin heavy chain (*MYH11* encoding smMHC) during SMC differentiation^27^. However, epigenetic, transcriptional and proteomic profiles of these cellular models are missing, which is critical to envision their use in the study of regulatory elements involved in the genetic risk for cardiodiseases where arterial remodeling is impaired.

In the current study, we implemented two existing methods to differentiate three iPSCs lines through the mesodermal lineage using either RepSox (R-SMCs) or TGF-β1 and PDGF-BB (TP-SMCs) during the second half of differentiation process^27^ (**Graphical Abstract**). We characterized chromatin accessibility and gene expression and searched for key transcriptional regulators at main steps of differentiation. We then examined the expression of genes involved in key SMC cellular processes in R-SMCs, TP-SMCs, as well as human primary coronary SMCs. Finally, we examined how cardiovascular disease associated variants were likely to be enriched in regulatory elements in these cell lines. Altogether, our results provide an important resource to evaluate iPSC-derived SMCs as potential cellular models for cardiovascular disease involving arterial remodeling.

## Results

### Cellular features of iPSCs-derived SMCs models from 2 differentiation protocols

We first characterized the cellular phenotypes of SMCs obtained using either RepSox (R-SMCs) or TGF-β/PDGF-BB (TP-SMCs) protocols. Cells differentiated using both protocols expressed several typical SMC markers, including SM α-actin, SM22 and calponin (Figure 1A). Cells treated with RepSox (R-SMCs) expressed in addition higher levels of smMHC (Figure 1A). Using cell viability and EdU incorporation assays we found that SMCs differentiated using TGF-β1 and PDGF-BB (TP-SMCs) had a higher rate of proliferation compared to R-SMCs (Figure1B-D). Assessment of migration capacity of cells using Transwell® (Figure 1E) and wound healing (Figure 1F-G) assays indicated a lower migration capacity for R-SMCs compared to TP-SMCs. Stimulation with PDGF-BB had limited impact on migration capacities of both SMC phenotypes (Figure 1E). To characterize the cell contractile ability, we measured the ability of cells to contract a collagen gel lattice (Figure 1H). 20h after seeding, R-SMCs induced a ∼55% reduction of gel area, which was partially impaired by incubation with 2,3-butanedione monoxime (BDM), an inhibitor of myosin-actin interactions^28^. TP-SMCs induced a higher contraction of gel area (∼80%), but BDM had little effect on gel contraction, suggesting this was independent from actin-myosin interactions. Altogether, these results suggest that R-SMCs present several features of contractile/quiescent SMCs, whereas TP-SMCs presented features of a synthetic and proliferative SMC phenotype.

**Figure 1.**
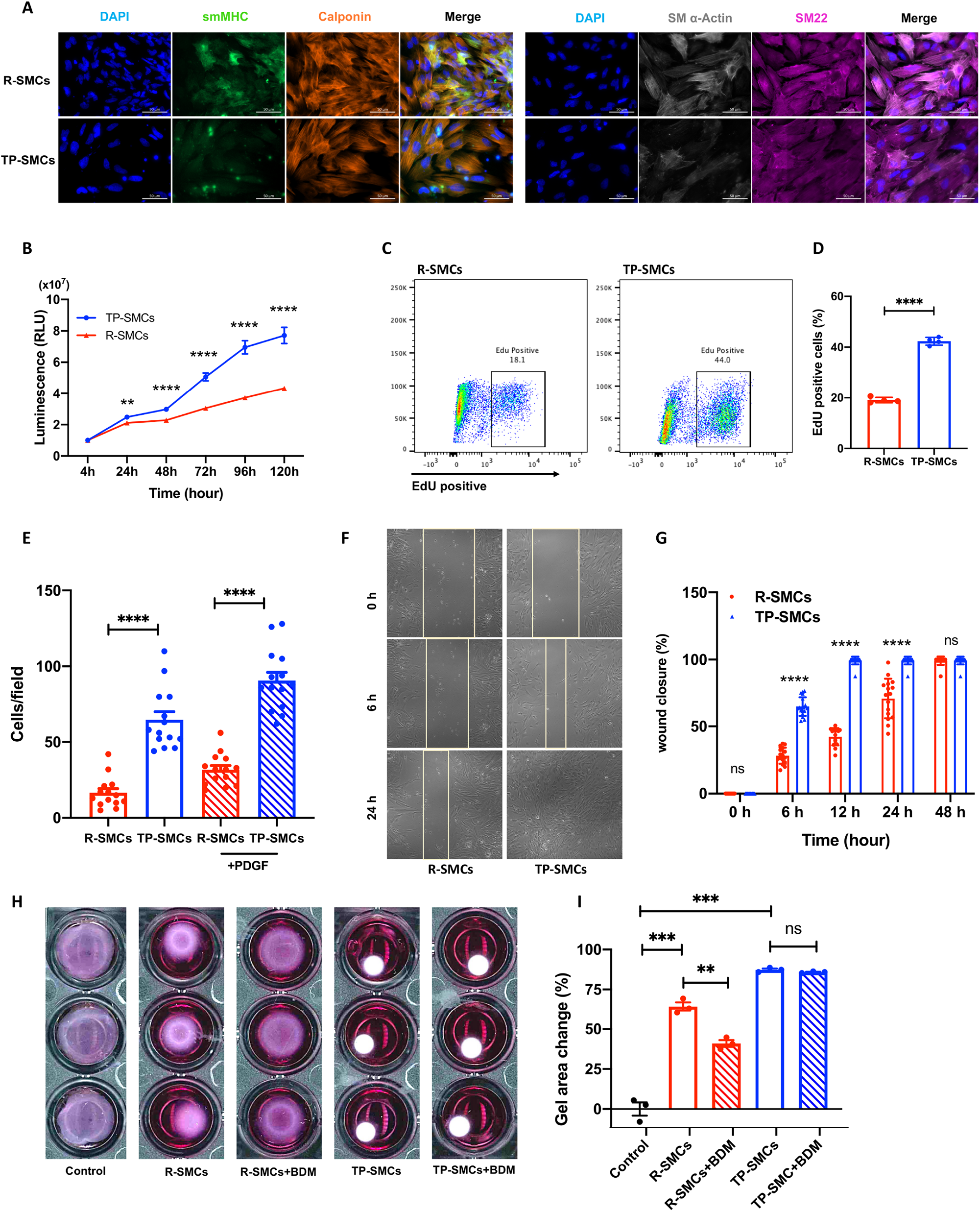
Characterization of iPSC derived R-SMCs and TP-SMCs. **A:** Representative immunofluorescence images of SMCs marker proteins smMHC (encoded by MYH11), Calponin (encoded by CNN1), α-Actin (encode by ACTA2) and SM22 (encoded by TAGLN) in iPSC derived R-SMCs and TP-SMCs. Scale bars: 50 µm. **B:** Measure of cell viability along a 120h culture period for R-SMCs (red) and TP-SMCs (blue). The same cell number were seeded at a low density. Cells were lysed and assayed using Cell Titer-Glo at 4h, 24h, 48h, 72h, 96h and 120h. Unadjusted P-value of comparison between sample groups using Student’s t-test is indicated: n=16, mean±SEM, **p<0.01, ****p<10^−4^, ns: not significant. **C**: Representative flow cytometry plots showing the EdU incorporation into cells in R-SMCs and TP-SMCs. **D:** Barplot representing the quantification of EdU positive cells. Unadjusted t-test: n=3, mean±SEM. Unadjusted t-test: ****p<10^−4^. **E:** Barplot representing mean±SEM of the number of cells per field counted following a Transwell assay assessing cell migration through polycarbonate membrane (8µm pore size, 20h culture) with or without PDGF. Unpaired t-test: n=14, mean±SEM, ****p<10^−4^. **F:** Representative images from scratch wound repair assay in R-SMCs, TP-SMCs at 0h, 6h, 24h. Delimited zone represents the wound area used for measurements. **G:** Barplot representing mean±SEM of the percentage of wound closure at 6h, 12h, 24h and 48h. Wound area was measured by Image J and compared to wound area measured at 0h. Unadjusted t-test: n=16, mean±SEM, ****p<10^−4^, ns: not significant. **H:** Representative images showing contraction of a collagen lattice by R-SMCs and TP-SMCs. **I:** Barplot representing mean±SEM of the percentage of gel area reduction after 20h for a collagen gel lattice seeded with no cells, or 2.5×10^5^ R-SMCs or TP-SMCs, with or without 10 mM 2,3-butanedione monoxime (BDM). Unadjusted t-test: n=3, mean±SEM, **p<0.01, ***p<0.001, ns: not significant.

### Global assessment of gene expression and chromatin accessibility profiles during iPSC differentiation into SMCs

To fully assess epigenetic features during differentiation, we initiated the differentiation of 3 independent iPSC lines (1 male, 2 females) into SMCs using the 2 protocols and generated polyA+-RNA-Seq and ATAC-Seq data at 6 time points of the differentiation process (day0, day3, day8, day12, day16 and day24) (Figure 2A). Hierarchical clustering of samples based on either RNA-Seq (Figure 2B) or ATAC-Seq (Figure 2C) data was overall consistent with the stage of differentiation process, with an early differentiation cluster of cells from day0 and day3, a mid-differentiation cluster including cells from day8 and day12 and a cluster of late differentiation stages (day16 and day24, Figure 2B-C). Interestingly, TP-SMCs at day 24 formed an additional separated cluster based on both RNA-Seq and ATAC-Seq results (Figure 2B-C). Overall, both gene expression and open chromatin maps efficiently discriminated iPSC to SMCs differentiation stages.

**Figure 2.**
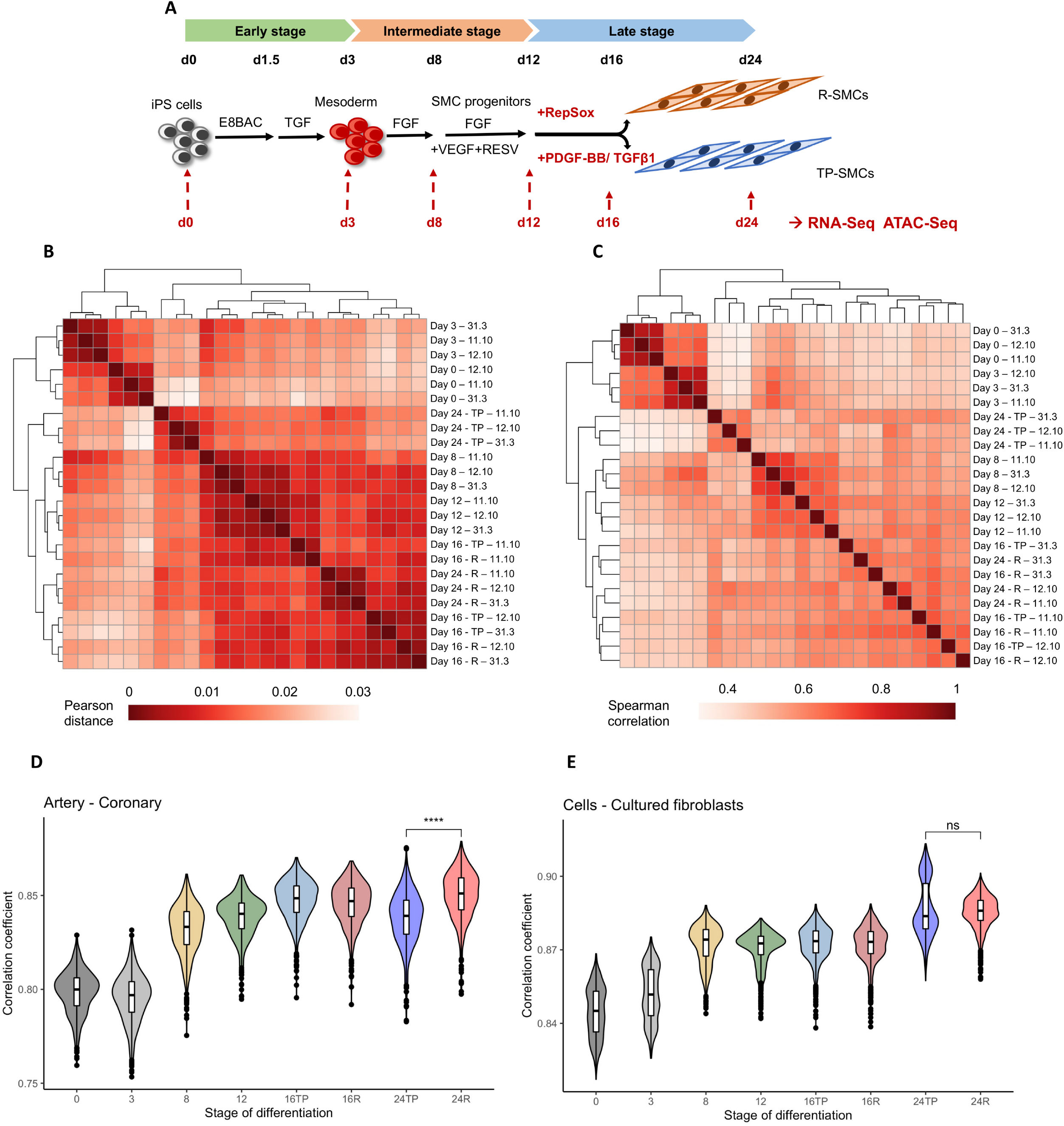
Global assessment of expression and chromatin accessibility profiles during differentiation. **A:** Illustration of the differentiation protocols and time points analysed using RNA-Seq and ATAC-Seq. **B:** Hierarchical clustering of samples based on Pearson distance between samples in RNA-Seq analysis of 3 hiPSC derived cell lines at 6 selected time points. **C:** Hierarchical clustering of samples based on Spearman correlation between samples in ATAC-Seq analysis of 3 hiPSC derived cell lines at 6 selected time points. **D-E:** Correlation of gene expression profiles between cells isolated at different stages during differentiation and gene expression profiles in coronary artery (D) and cultured fibroblasts (E) samples from GTEx database. Unadjusted Wilcoxon test: ****p<10^−4^, ns: not significant (p>0.05).

We leveraged existing expression data to assess putative similarities in terms of gene expression between iPSC-derived SMCs and GTEx cultured cells and tissues. To this aim, we calculated the global correlation between gene expression profiles of iPSC-derived cells and tissues. Gene expression profiles from iPSCs-derived SMCs were highly correlated with expression profiles from fibroblasts and SMC-rich tissues, including arteries (Supplementary Figure 1A). We found that gene expression correlations with fibroblasts and artery tissue increased during the differentiation process to reach a maximum at the end of the process on day24 (Figure 2D-E). Cells differentiated using RepSox presented a higher correlation with the expression profiles of coronary artery tissue (Figure 2D) as well as other artery tissues (Supplementary Figure 1B-C), whereas both protocols led to similar levels of correlation with the gene expression profiles of cultured fibroblasts (Figure 2E).

### Identification of key drivers of iPSCs differentiation to SMCs

To identify the main transcriptional and regulatory events occurring during differentiation, we compared gene expression and chromatin accessibility profiles between successive time points, focusing on the transitions representing most of the variability in our datasets, namely early (day 0 to day 3), intermediate (day 3 to day 8) and late (day 12 to day 24) stages, with a special focus on differences between RepSox and TGF-β1/PDGF-BB protocols (Figure 2A).

Early-stage differentiation was characterized by an upregulation of genes related to pattern specification and regionalization, with “mesenchyme” (*P*=4×10^−26^) and “mesoderm development” (*P*=5×10^−14^) among the most enriched gene ontology (GO) terms, concomitant with a downregulation of pluripotency factors such as *SOX2* (Figure 3A, Supplementary Figure 2, Supplementary files 1-2). At the intermediate stage, upregulated genes were markedly related to GO terms “extracellular matrix organization” (*P*=1×10^−50^), “connective tissue development” (*P*=2×10^−21^) and “regulation of blood circulation” (*P*=5×10^−13^), while factors associated with early mesoderm development such as *ZIC2* and *ZIC3* were downregulated^29^ (Supplementary file, Supplementary Figure 3, Supplementary files 1-2). Markers for endothelial cells such as *CDH5* and *KDR* were transiently expressed at this stage together with mesenchymal genes such as *SNAI1* indicating a persistent multipotent state of cells at this stage of differentiation (Figure 3A).

**Figure 3.**
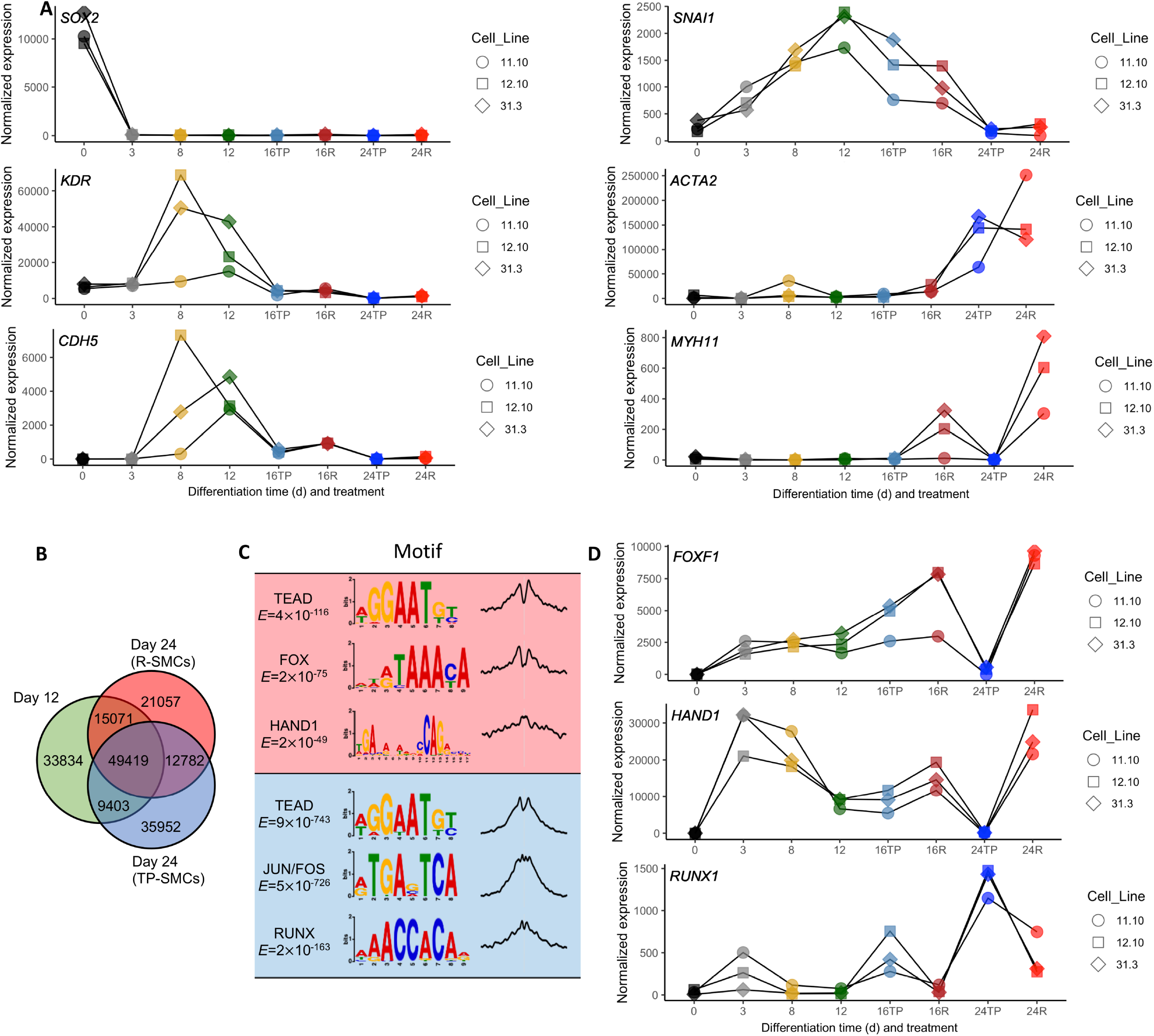
Key markers and drivers of differentiation. **A:** Normalized gene expression of *SOX2, SNAI1, KDR, ACTA2, CDH5, MYH11* during the differentiation of 3 iPSC cell lines. **B:** Venn diagram representation of overlaps between open chromatin regions of cells at day 12 and day 24 of differentiation (either R-SMCs or TP-SMCs). **C:** Most enriched transcription factors motifs in peaks specific to R-SMCs (upper panel) and TP-SMCs (lower panel). Motif logo, *E*-value and central enrichment with respect to peak summits were obtained using MEME-ChIP software. **D:** Normalized gene expression of *FOXF1, HAND1* and *RUNX1* during the differentiation of 3 iPSC cell lines.

During late differentiation, genes involved in “muscle contraction” (*P*=2×10^−10^) and “vascular process in circulatory system” (*P*=2×10^−9^) emerged as enriched in upregulated genes in R-SMCs. On the other hand, genes involved in “endoplasmic reticulum stress response” (*P*=2×10^−^ _13_), macroautophagy (*P*=4×10^−9^), and “protein secretion” (*P*=3×10^−6^) were the most enriched terms among upregulated genes in TP-SMCs (Supplementary file 1). Open chromatin regions specific to R-SMCs were enriched in binding sites for Forkhead-Box (FOX) and HAND1 transcription factors, whereas AP-1 and RUNX binding sites were relatively enriched in open chromatin regions specific to TP-SMCs (Figure 3B-C, Supplementary file 2). This was consistent in particular with the expression patterns of *FOXF1, HAND1* and *RUNX1* transcription factors (Figure 3D).

### RepSox differentiation induces the expression of markers related to contractile signaling

We compared transcriptomic profiles at day 24 of differentiation of both protocols with primary human coronary artery SMCs (Primary-SMCs) and ES-derived SMCs using an alternative method involving treatment with BMP4 and Wnt3a through a 22-day protocol (BW-SMCs)^30^. All protocols led to high levels of expression of SMCs markers SM α-actin (*ACTA2*) and calponin 1 (*CNN1*) genes. However, only R-SMCs presented higher levels of expression of *MYH11*, an established marker for smooth muscle contraction (Figure 4A). Of note, we found higher expression of transgelin gene (*TAGLN*, encoding the SMC marker SM22) in TP-SMCs, compared to the other iPSC-derived SMCs (Figure 4A). Hierarchical clustering of RNA-Seq datasets based on Pearson distance between samples showed that R-SMCs, TP-SMCs, primary SMCs and BW-SMCs formed separate clusters, confirming they represent distinct SMC phenotypes (Figure 4B). Interestingly, TP-SMCs showed more similarity of gene expression patterns with primary SMCs, compared to the other iPSC-derived SMCs, possibly reflecting the synthetic transition of primary SMCs in cell culture^31^. Principal component analyses provided consistent findings with Pearson correlation analyses (Figure 4B, Supplementary Figure 4).

**Figure 4.**
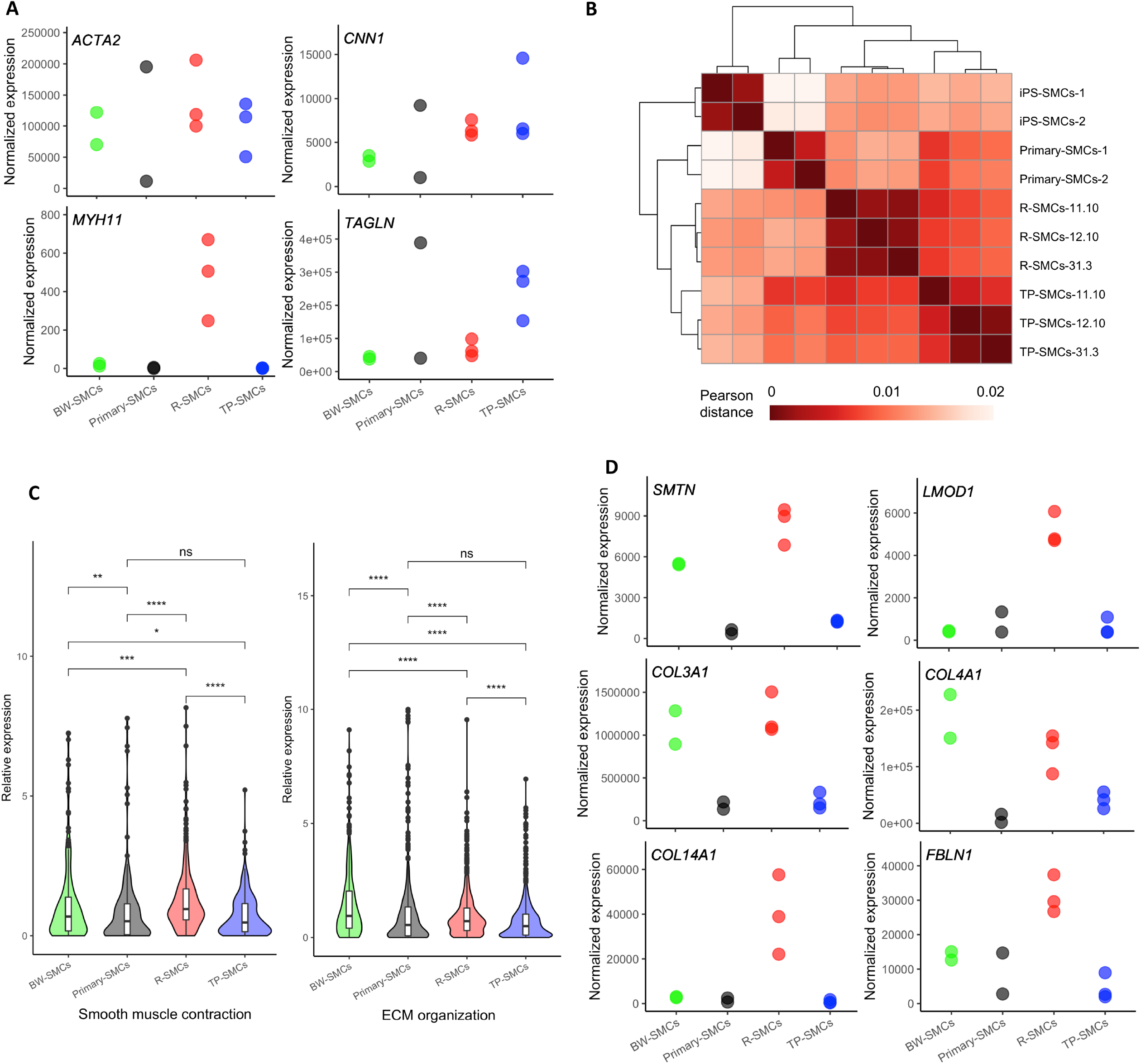
Comparison of gene expression profiles in four SMC models. **A:** Normalized gene expression of SMC markers gene *ACTA2, CNN1, MYH11* and *TAGLN* in BW-SMCs, R-SMCs, TP-SMCs and primary SMCs. **B:** Hierarchical clustering of samples based on Pearson distance between samples in RNA-Seq analysis of in BW-SMCs, R-SMCs, TP-SMCs and primary SMCs. **C:** Violin plot indicating the relative expression distribution of genes associated with gene ontology term “smooth muscle contraction” (left panel) and “Extracellular matrix (ECM) organization” (right panel) in BW-SMCs, R-SMCs, TP-SMCs and primary SMCs. Unadjusted *P*-value of comparison between sample groups using Wilcoxon test is indicated: *p<0.05, *p<0.01, ***p<0.001, ****p<10^−4^, ns: not significant. **D:** Normalized gene expression of SMC markers genes *SMTN* and *LMOD1*, and extracellular matrix genes *COL3A1, COL4A1, COL14A1* and *FBLN1* in BW-SMCs, R-SMCs, TP-SMCs and primary SMCs.

We calculated the relative expression of genes associated to Gene Ontology processes related to SMCs biology in R-SMCs, TP-SMCS, BW-SMCs and Primary-SMCs. We found that R-SMCs presented an overall higher expression of genes annotated as involved in “Smooth muscle contraction” compared to TP-SMCs (*P*_*adj*_= 3×10^−13^), Primary-SMCs (*P*_*adj*_= 1×10^−9^) and BW-SMCs (*P*_*adj*_= 1×10^−3^) (Figure 4C). Particularly, we observed higher expression of genes regulating the assembly of SMC contractile fibers, such as smoothelin (*SMTN)* and leiomodin 1 (*LMOD1)* (Figure 4D). R-SMCs also expressed more genes involved in the regulation of VSMCs differentiation, smooth muscle tissue development and SMC proliferation, although in comparable levels to primary SMCs or BW-SMCs (Supplementary Figure 5). Unexpectedly, R-SMCs expressed higher levels of genes involved in “extracellular matrix organization”, compared to TP-SMCs (*P*_*adj*_= 3×10^−10^) and primary-SMCs (*P*_*adj*_= 2×10^−4^) (Figure 4C). This involved in particular several collagen subtypes, including fibrillar (e.g *COL1A1, COL3A1*, and *COL5A1*) and nonfibrillar (e.g *COL4A1* and *COL14A1*) collagens, and fibulin 1 (*FBLN1*), a component of fibronectin-containing matrix fibers (Figure 4D, Supplementary Figure 6). Both TP-SMCs and R-SMCs expressed relatively low levels of fibronectin (*FN1*), elastin (*ELN*), and fibrillin 1 (*FBN1*), which are essential components of the arterial extracellular matrix (Supplementary Figure 6).

### Analysis of extracellular matrix generated by iPSCs-derived SMCs

Production and maturation of extracellular matrix requires multiple steps beyond gene expression. To characterize the production of extracellular matrix (ECM) by R-SMCs and TP-SMCs, we purified ECMs from decellularized cultures of R-SMCs, TP-SMCs and Primary-SMCs, and analyzed their composition using mass spectrometry and label-free quantification using MaxQuant (Figure 5A). We quantified from 200 to more than 1100 different proteins per samples (Supplementary Figure 7). Overall, the composition of ECMs were similar between R-SMCs and TP-SMCs, although more proteins were detected from R-SMCs-derived ECM extracts (Supplementary Figure 7A-B), with fibronectin 1, tenascin C, type I and type IV collagens as the most frequently detected ECM constituents (Supplementary Figure S7B, Supplementary file 3). Conversely, several differences were observed between ECMs generated by R-SMCs and TP-SMCs. In particular, several proteins required for basement membrane assembly such as peroxidasin, involved in crosslinking and bundling of collagen IV fibrils^32^, and laminins, particularly laminin subunit gamma 1 (LAMC1), were more present in R-SMCs-generated ECM (Figure 5B, Supplementary Figure 7B). Among most enriched proteins in R-SMCs-ECM, we highlight thrombospondin 1 (THBS1) and cell surface glycoprotein MUC18 (MCAM), involved in the regulation of cell adhesion and migration (Figure 5B). In TP-SMCs-ECM, we found enrichment for EMILIN1, a regulator of elastic fibers deposition and inhibitor of TGF-β signaling^33^, thrombospondin type-1 domain-containing protein 4 (THSD4), involved in fibrillin 1 deposition, plasminogen activator inhibitor 1 (SERPINE1) and periostin (POSTN), both key regulators of wound repair pathways^34, 35^. Differential detection of proteins in the ECM was highly correlated with differential gene expression (R=0.33, P=8×10^−20^), showing that most of the observed differences are likely to stem from changes in the cell regulatory profiles (Figure 5C).

**Figure 5.**
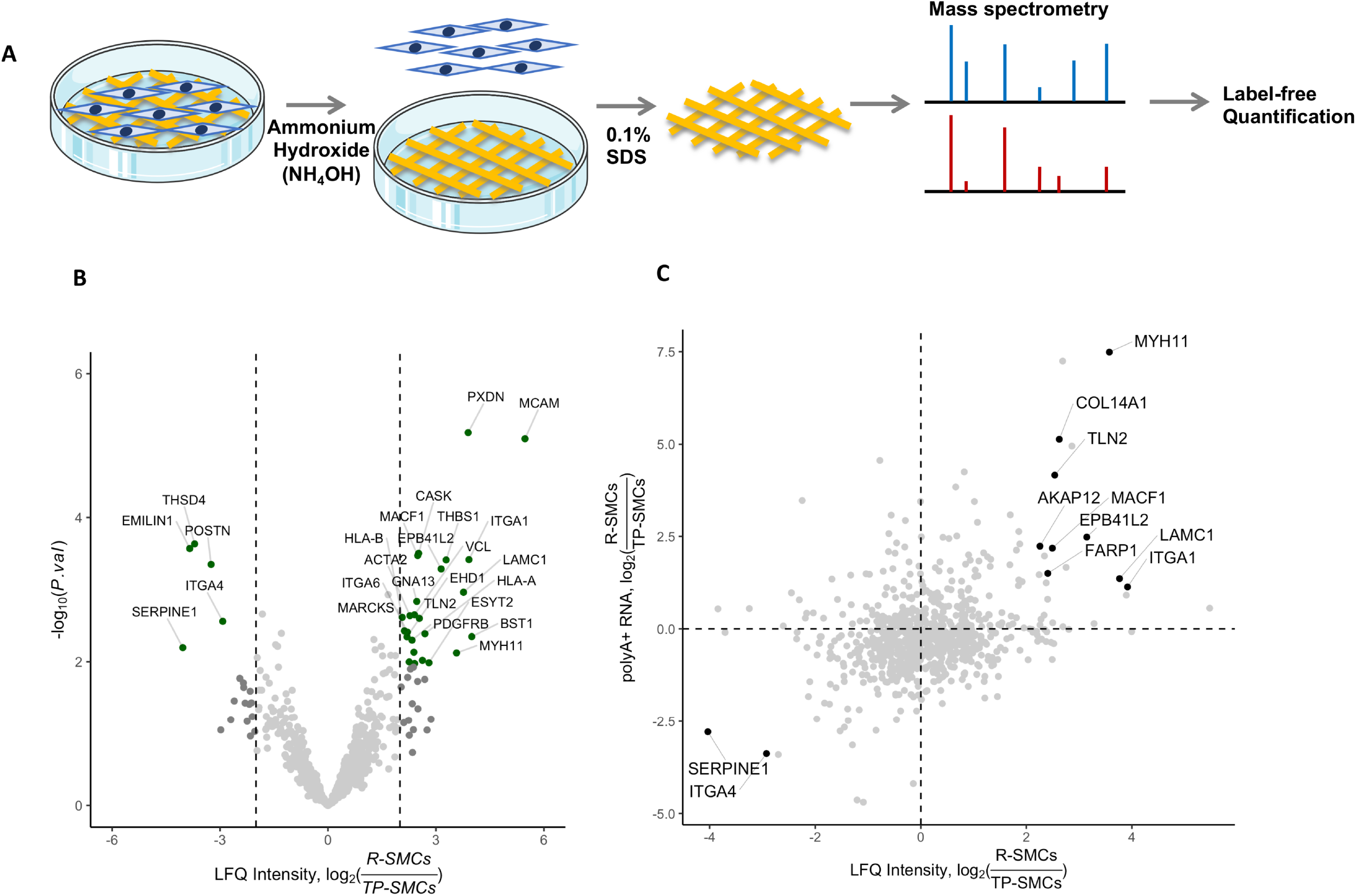
Analysis of ECM synthesis by iPSCs-derived SMCs and primary SMCs. **A:** Illustration of the analysis of ECM generated by R-SMCs, TP-SMCs and primary-SMCs using mass spectrometry on decellularized ECM extracts. **B:** Volcano plot representation of differentially detected proteins between extracellular extracts of TP-SMCs and R-SMCs. x-axis represents the log2 Fold Change of label-free quantification (LFQ) intensities, while y-axis represents P-value on a negative log10 scale. Proteins with FDR < 0.2 and absolute log2 fold change > 2 are colored in green **E:** Scatter plot representation of RNA differential expression (y-axis, log2 Fold Change) vs extracellular protein differential detection (x-axis, log2 Fold Change). Genes/protein with FDR < 0.2 for both differential expression and differential extracellular protein detection analyses are highlighted.

### Relevance of R-SMCs and TP-SMCs to study risk variants for cardiovascular disease

The genetic basis of complex cardiovascular diseases is mostly impacted by non-coding variants located in specific regulatory regions. To assess iPSC-derived SMCs as cellular models to study these variants, we compared their open chromatin regions (OCRs) to those of primary SMCs derived from human coronary and carotid arteries, in addition to whole healthy coronary arteries^16, 36^. Most OCR identified in arteries were found of at least one cellular model, while a large number of OCR (>83,000) were specific to primary cultured SMCs (Figure 6A). About 37,000 OCRs were shared among all tissues and cells and were mainly located in gene promoters. The remaining groups of OCRs were more evenly located in promoters, introns and intergenic regions (Supplementary Figure 8).

**Figure 6.**
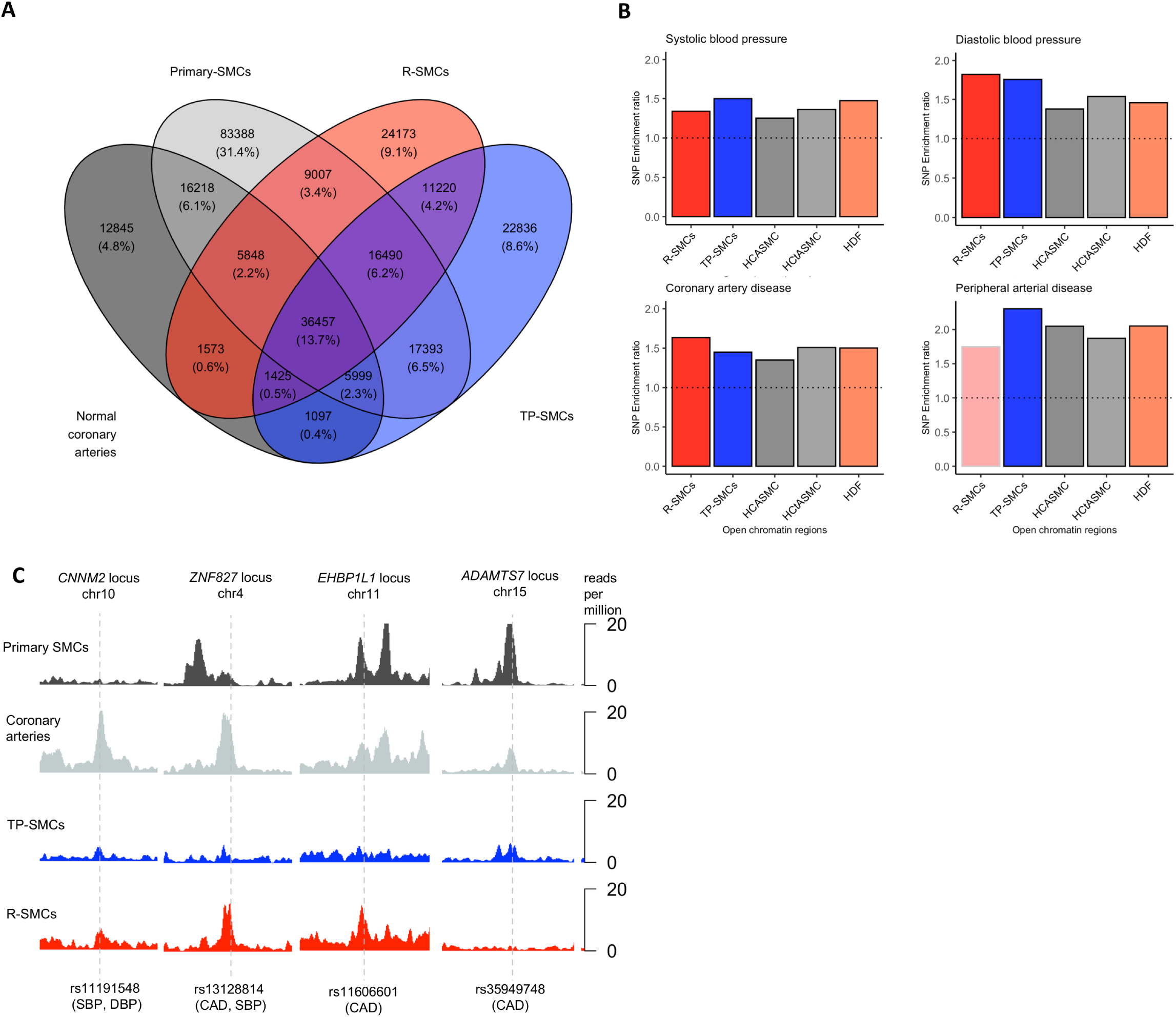
Assessment of iPS-derived SMCs as cellular models for arterial diseases. **A:** Venn diagram representing the overlaps between peaks detected in in ATAC-Seq analysis of normal coronary arteries, R-SMCs, TP-SMCs and primary human carotid (HCtASMCs) and coronary (HCASMCs) artery SMCs. **B:** Enrichment (Fold ratio) of disease associated variants in peaks identified in ATAC-Seq analysis of R-SMCs, TP-SMCs, coronary artery SMCs (HCASMC), carotid artery SMCs (HCtASMC) and dermal fibroblasts (HDF). Black outline represents conditions with significant enrichment (*P*.*adj* <0.05). **C**: Genome browser visualization of chromatin accessibility at proximity of 4 example variants associated to regulation of systolic blood pressure (SBP), diastolic blood pressure (DBP) and/or coronary artery disease (CAD).

We then computed the enrichment for common genetic variants associated with several cardiovascular diseases and traits in the OCRs of R-SMCs, TP-SMCs, primary SMCs and dermal fibroblasts. We found a significant enrichment of associated variants in OCRs of R-SMCs, TP-SMCs and primary cells for variants associated to systolic and diastolic blood pressure (Figure 6B). Similarly, R-SMCs and TP-SMCs-OCRs were enriched in variants associated to coronary artery disease (Figure 6B). We found an enrichment variant associated to peripheral arterial disease only TP-SMCs-OCRs (Figure 6B), while variants associated to intracranial aneurysm were enriched in R-SMCs-OCRs only (Supplementary Figure 9). We did not find a significant enrichment in variants associated to ischemic stroke, abdominal aortic aneurysm and migraine in both iPSCs-derived SMCs (Supplementary Figure 9). At the loci level, we found that candidate variants for causality overlapped accessible regulatory elements in some cases in one specific cell culture condition. This was for example the case for rs11191548, a likely causal variant associated to blood pressure located in *CNNM2* 3’UTR on chromosome 10, which is accessible in coronary arteries, R-SMCs and TP-SMCs but not in primary SMCs (Figure 6E). Similarly, rs1318814, a variant associated to CAD and blood pressure traits, intronic to *ZNF827* on chromosome 4, overlapped with a regulatory element mostly activated in coronary artery and R-SMCs, but not in TP-SMCs (Figure 6E). Another CAD variant, rs35949748 intronic to *ADAMTS7* on chromosome 15 was accessible in TP-SMCs but not in R-SMCs (Figure 6E).

### Vascular tone related signaling is increased in RepSox-derived SMCs

To determine whether the differences at the level of chromatin accessibility were also observed at the gene expression level, we analyzed the relative gene expression in SMCs cellular models of genes annotated as involved in cardiovascular diseases in the Disgenet database^37^. Globally, no major differences were observed between cell culture conditions for most diseases (Supplementary file 3). However, we found that the majority of genes annotated as involved in hypertension were more expressed in R-SMCs and BW-SMCs, compared to TP-SMCs or primary SMCs (Figure 7A). We also found a stronger expression of genes involved in coronary artery disease in R-SMCs, compared to TP-SMCs (Supplementary Figure 9). Among the genes with stronger differential expressions, we highlight the receptors for angiotensin and endothelin contractile signals (*AGTR1* and *EDNRA*, Figure 7B), as well as several genes with documented effect in SMCs phenotypic modulation in coronary artery disease such as *COL4A1*^38^, *LMOD1*^39^ (Figure 4D), *TCF21*^*40*^ and *ADAMTS7*^41^ (Figure 7B), which were all strongly upregulated in R-SMCs (Supplementary file 1).

**Figure 7.**
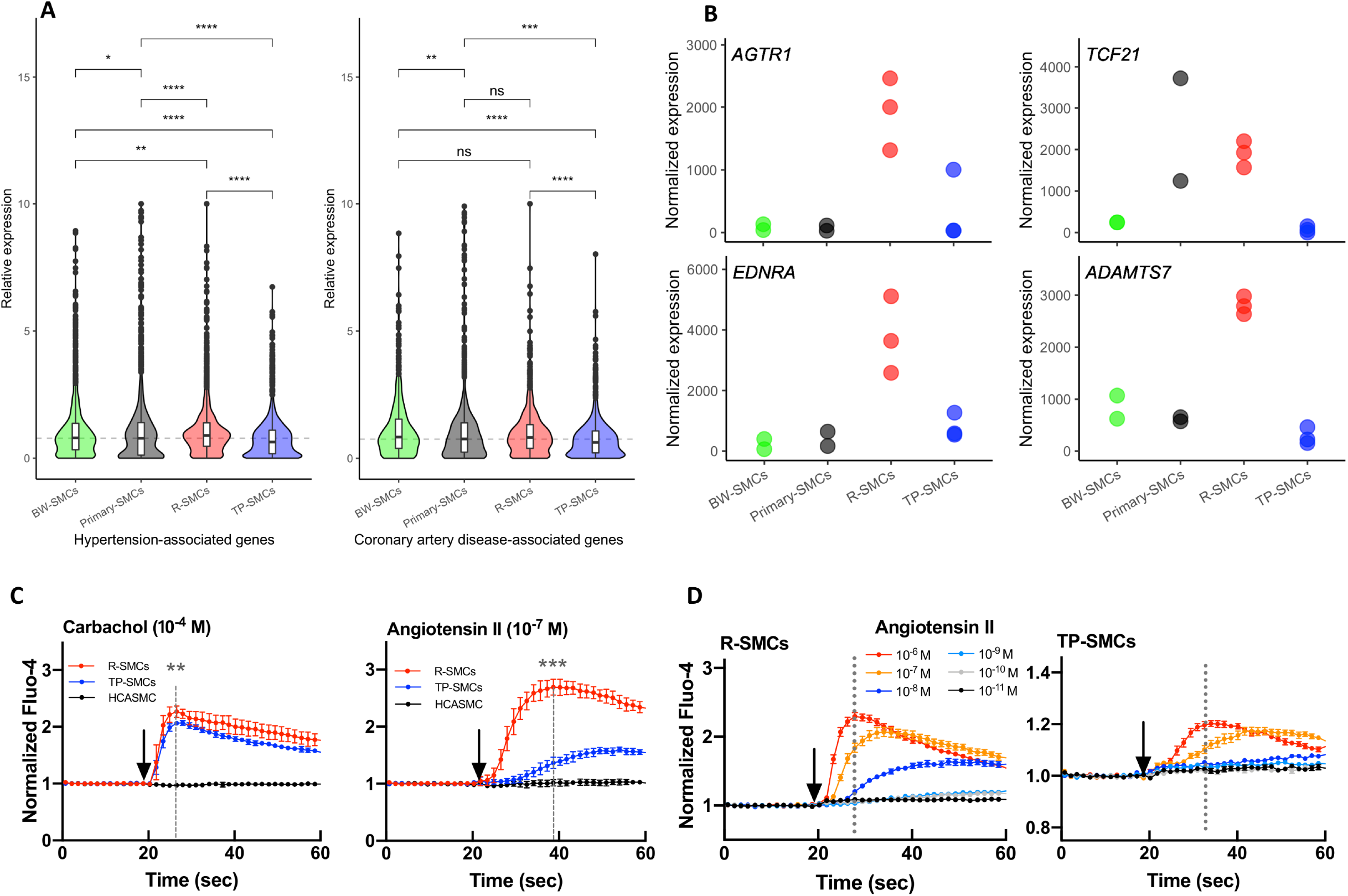
Blood pressure related signaling in iPSC-derived SMCs. **A:** Violin plot indicating the relative expression distribution of genes associated with hypertension and coronary artery disease in Disgenet database (Gene-disease association score > 0.1). Unadjusted *P*-value of comparison between sample groups using Wilcoxon test is indicated: *p<0.05, **p<0.01, **p<0.001, ***p<0.0001, ns: not significant. **B:** Normalized gene expression of hypertension associated genes *AGTR1* and *EDNRA* and coronary artery disease-associated genes *TCF21 and ADAMTS7* in BW-SMCs, R-SMCs, TP-SMCs and primary SMCs. **C**: Measure of fluorescence in cells labeled with intracellular calcium probe fluo-4 and treated by carbachol or Angiotensin II as indicated in R-SMCs (red), TP-SMCs (blue) and primary SMCs (black). Comparison of the fluorescence values (Student T-test) were performed at a maximum calcium signals for R-SMCs (dashed line): n=6, mean±SEM, **p<0.01, ***p<0.001. Arrows indicate the time points where the inducers were added to cells. **D:** Measure of fluorescence in R-SMCs or TP-SMCs labeled with intracellular calcium probe fluo-4 and treated by increasing doses of Angiotensin II. Dashed lines indicate the time points where maximum values were obtained. Arrows indicate the time points where Angiotensin II was added to cells.

To further define the response to contractile signals in iPSC-derived SMCs, we assessed the intracellular Ca^2+^ levels following treatment with either carbachol, an activator of muscarinic receptors or angiotensin II using fluo-4 fluorescent probes. Carbachol induced a rapid release of calcium both in R-SMCs and TP-SMCs, with a slightly higher peak value in R-SMCs, but no calcium release in primary SMCs (Figure 7C). Angiotensin II induced a rapid increase of calcium efflux and peaked at 15 seconds post treatment in R-SMCs (Figure 7C). By contrast, the calcium level of TP-SMC increased later and peaked at approximately 35 seconds post treatment (Figure 7D). No increase in fluorescence was detected in primary coronary artery SMCs. Also, R-SMCs were sensitive to a minimum angiotensin II concentration of 10^−8^ M, while TP-SMCs had no reaction at this concentration, indicating that R-SMCs had higher sensitivity to angiotensin II (Figure 7D).

## Discussion

In this study, we performed deep cellular phenotyping, epigenomic and proteomic depiction of SMCs derived from induced pluripotent stem cells (iPSCs) and compared them to existing data of primary SMCs and artery tissue. We found that differentiation using TGF-β and PDGF-BB led to a SMC phenotype similar to primary SMCs cultured from coronary arteries. On the other hand, differentiation with RepSox, a TGF-β inhibitor, led to a global increase of the expression of contractile markers, enhanced production of arterial extracellular matrix components, lower cellular proliferation and migration and higher correlation with arterial gene expression profiles. These cells presented phenotypic and transcriptomic similarities with contractile, quiescent phenotype of arterial SMCs. We found high potential for these models to the study of complex regulatory mechanisms at genetic risk loci involved in several arterial diseases, and that RepSox-derived SMCs may be particularly suitable to study pathways involved in blood pressure homeostasis.

A wide variety of methods to generate functional SMCs from pluripotent cells have been previously described with limited cellular or genomic characterization^20^. Here we provided an epigenomic characterization of two SMC differentiation protocols based on a shared initial framework inducing mesodermal differentiation from iPSCs^27^. While TGF-β and PDGF-BB are growth factors commonly used to induce SMC differentiation^23^, RepSox was identified using a screening approach aiming at maximizing the expression of *MYH11* gene^27^. RepSox functions as a selective TGFβR-1/ALK5 inhibitor which inhibits ALK5 autophosphorylation^42^ and was used to reprogramming of fibroblasts into iPSCs by replacing cMyc and Sox2^43^. Recently, it was reported to induce brown adipogenesis^44^ and promote the differentiation of sheep fibroblasts into adipocytes^45^. RepSox induction of contractile markers in SMCs was found to depend on activation of NOTCH signaling pathway^27^. However, we found that transcription factor binding sites for forkhead-box transcription factors and for HAND1 were highly enriched in chromatin regions specifically accessible in late differentiation using RepSox. *FOXF1*, in particular, was highly upregulated by RepSox while it was silenced using TGF-β and PDGF-BB. *Foxf1* is essential to early vascular development in mice and is also involved in visceral smooth muscle development but was not found to be expressed in arterial smooth muscle^46, 47^. Other forkhead family transcription factors such as FOXO3A and FOXM1 are known to regulation the differentiation and maintenance of arterial SMCs^48, 49^. The overexpression of FOXF1 in response to RepSox is likely to compensate for other forkhead box factors expressed at higher levels in artery tissue, or during later developmental stages. Conversely, differentiation of SMCs using TGF-β and PDGF-BB appears to specifically activate RUNX-dependent transcription. *RUNX1* is known to promote extracellular matrix formation^50^, and was identified as involved in myofibroblast differentiation and proliferation in various contexts^51-53^.

Our study provided several lines of evidence supporting iPSC derived SMCs using RepSox present many features of contractile phenotype-like SMCs. Compared to TP-SMCs, R-SMCs displayed lower proliferation, lower migration, capacity to contract a collagen matrix in an actin-myosin dependent way and higher calcium release capacity in response to contractile signals. Overall, the expression of genes involved in the regulation of smooth muscle contraction was higher in R-SMCs compared to all the other SMCs tested. The regulation of SMC contractility plays a key role in several arterial diseases including aortic aneurysm and dissection, atherosclerosis and hypertension^54-57^. We found that genes associated to hypertension and coronary artery disease were expressed at higher levels in R-SMCs than in TP-SMCs or cultured primary SMCs and that genetic variants associated to coronary artery disease and blood pressure regulation were highly enriched in the open chromatin regions of R-SMCs. These cells showed high sensitivity to angiotensin II, an important mediator of blood pressure regulation, arterial ageing and aneurysm^58^. Altogether, these results point towards R-SMCs as a very promising cellular model for functional studies on the role of SMCs in arterial diseases involving impaired vascular tone.

## Limitations

Our work presents several limitations. First, we deeply studied two SMC differentiation protocols, but many other protocols, involving for example different growth factors or embryoid bodies, were proposed and may represent highly useful approaches that we could not evaluate in this work. Second, these two protocols are based on mesoderm lineage and it was previously shown that embryonic origin may largely affect the phenotype of SMCs^18, 20^. Third, although we tested the differentiation of SMCs using three fully independent cell lines, this was not sufficient to estimate the variability due to different iPSC clones, and in particular we could not test the effect of cell sex on the obtained SMC phenotypes.

## Conclusion

Our results provide a valuable resource to evaluate different potential SMC differentiation protocols to study cardiovascular disease genetic loci. We provide integrative evidence based on gene expression regulation, secretome and cellular phenotyping supporting RepSox-derived SMCs as a promising model to study SMC contractile function in relation to arterial diseases, specifically in relation to vascular tone regulation.

## Methods

### iPSC culture

Human iPSC line SKiPS-31.3 was by reprogramming of human dermal fibroblast of a healthy male adult volunteer as previously described^59^. iPS cell lines 11.10 and 12.10 were purchased from Cell Applications (San Diego, CA) and were identified as female using *SRY* PCR. iPSC-clones C1-C4 used for mass spectrometry analyses were derived from iPSC 11.10 following transfection with pSpCas9(BB)-2A-Puro without guide RNA^60^, treatment for 48h with puromycin, colony isolation and assessment of normal karyotype and expression of pluripotent markers. All iPSC lines were maintained in mTeSR™ Plus medium (STEMCell Technologies) changed every other day. The iPSCs were passaged every week by scraping cells using tips. Cultured plates were coated with Matrigel (Corning, Corning, NY, USA).

### iPSC differentiation

Base differentiation medium (E5 medium) contained L-ascorbic acid-2-phosphate magnesium (64 mg/l), sodium selenium (14 *μ*g/l), NaHCO_3_ (543 mg/l), transferrin (10.7 mg/l) added to a DMEM/F12 base medium (Thermo Fisher Scientific, Waltham, MA, USA), as previously described^61^. At first day of differentiation (day 0), iPSCs were dissociated into single cells by Accutase (Thermo Fisher Scientific) for 8 mins at 37 °C. Cells were seeded on Matrigel-coated dish at 10^5^ cells/cm^2^ and cultured in E8BAC medium (E5 medium plus 5 ng/ml BMP4, 25 ng/ml Activin A, 19.4 mg/l insulin, 10 *μ*M Y27632 and 1 *μ*M CHIR99021) for 36 hours (Day2). Then, cells were dissociated by Accutase and seeded on a new dish coating Matrigel at 1.6 × 10^4^ cells/cm^2^ in E6T medium (E5 medium supplemented with 19.4 mg/l insulin, 1.7 ng/ml TGF-*β*1 and 10 *μ*M Y27632) for 18 hours. Cells would achieve 100% confluence on day 6 for the optimal differentiation efficiency. From day 3 to day 7, cells were cultured with E5F medium (E5 medium supplemented with 19.4 mg/l insulin and 100 ng/ml FGF2). The E5F medium was changed at day 3, day 4 and day6. From day 8 to day 11, the cells were treated with FVR medium (E5 medium supplemented with 19.4 mg/l insulin, 50 ng/ml VEGF and 5 *μ*M RESV) and changed every other day. From day 12, cells started to differentiate into SMC phenotypes. For TP protocol, cells were cultured in E6-TP medium (E5 medium supplemented with 19.4 mg/l insulin, 5 *μ*M RESV, 10 ng/ml PDGFbb and 1.7 ng/ml TGF-*β*1). For RepSox protocol, cells were treated with E6-R medium (E5 medium supplemented with 19.4 mg/l insulin, 5 *μ*M RESV, 25 *μ*M Repsox). All media were changed every other day after day 12. On day 16, cells were cryopreserved when necessary or split in a new Matrigel-coated dish at 1×10^5^ cells/cm^2^ cell density. The volume of medium was increased to 1.5x volume over standard volume for day 1 and day 8-24. After day 24, R-SMCs were maintained in E6R medium, TP-SMCs and primary SMCs were grown in DMEM supplemented with 5 *μ*g/ml insulin, 0.5 ng/ml EGF, 2 ng/ml FGF and 5% FBS.

### RNA preparation

During the differentiation process, cells were collected at day 0 (iPSCs), day 3, day 8, day 12, day 16, and day 24. From iPSCs to day 12, cells were dissociated with Accutase for 8 mins at 37 °C and washed with PBS once. From day 16 to day 24, cells were split with 1 mg/ml collagenase IV plus 0.25 mg/ml Dispase for 10 mins, TryPLE for 10 mins and then washed once with PBS. The purification of total RNA was performed by the RNeasy Plus Mini kit (Qiagen, Hilden, Germany) following instructions of the manufacturer.

### RNA-Seq

Isolated total RNAs were quantified using Qubit ds DNA HS assay kit. PolyA+ mRNA libraries were prepared from 200ng-1µg of total RNA using QIAseq Stranded RNA library preparation kit (Qiagen) according to manufacturer’s instruction. Libraries were sequenced on a NextSeq500 instrument (Illumina, San diego, CA, USA) using NextSeq 500/550 High Output Kit v2 (75 cycles). Reads were demultiplexed using bcl2fastq2 (v2.18.12) and adapters were trimmed using Cutadapt (v1.15). Reads were then mapped to human genome (GRCh38.104) using STAR aligner (v2.7.9a) with following options: --outFilterType BySJout -- outFilterMultimapNmax 20 --alignSJoverhangMin 8 --alignSJDBoverhangMin 1 -- outFilterMismatchNmax 999 --outFilterMismatchNoverReadLmax 0.04 --alignIntronMin 20 - -alignIntronMax 1000000 --alignMatesGapMax 1000000. For BW-SMCs, raw reads were retrieved from ENCODE database experiment ENCSR236URT (two libraries from isogenic clones) and analyzed using same pipeline. We used per gene read counts as direct input for differential expression analysis using DESeq2 (v1.32.0) package in R (v4.1.0), keeping only genes with mean read count over 1^62^. We transformed the count matrix using variance stabilizing transformation with option blind=FALSE. Differentially expressed genes were determined using res function and log2 fold changes were determined using lfcShrink function with ashr shrinkage estimator^63^. We calculated Pearson distance between samples and performed clustering using Dist and pheatmap functions. Normalized gene expression values were retrieved using the count(normalized=TRUE) function in DESeq2. Relative gene expression was calculated as the ratio of normalized expression over the mean of normalized expressions in all presented conditions. Gene ontology term enrichments were calculated using clusterProfiler package (v4.0.5). To compare our RNA-Seq data with expression data from GTEx consortium, we took for each tissue/sample the floor of median gene expression value in reads per 100 million reads as input to DESeq2^64^. We transformed the count matrix using variance stabilizing transformation with option blind set to FALSE and performed correlation using cor function. To display relative gene expression by gene ontology terms, full list of human genes annotated to terms “regulation of smooth muscle contraction” (GO:0006940), “smooth muscle cell proliferation” (GO:0048659), “smooth muscle cell migration” (GO:0014909), “smooth muscle tissue development” (GO:0048745), “regulation of vascular associated smooth muscle cell differentiation” (GO:1905063) and “extracellular matrix organization” (GO:0030198) were retrieved from Gene Ontology webserver (http://amigo.geneontology.org/) on January 21^st^ 2022^65^. To display relative gene expression for disease associated genes, full list of disease associated genes were retrieved from Disgenet database using disease2gene function from disgenet2r package (v0.99.2), with options database set to ‘ALL’ and score between 0.1 and 1^37^. The following diseases were included: Myocardial Infarction (C0027051), Coronary Arteriosclerosis (C0010054), Coronary Artery Disease (C1956346), Atherosclerosis (C0004153), Cerebral Hemorrhage (C2937358), Aortic Diseases (C0003493), Vascular Diseases (C0042373), Hypertension (C0020538), Carotid Artery Diseases (C0007273), Cerebrovascular accident (C0038454), Intracranial Hemorrhage (C0151699), Cerebral Small Vessel Diseases (C2733158), Arterial Occlusive Diseases (C0003838), Peripheral Vascular Diseases (C0085096), Peripheral Arterial Diseases (C1704436), Aortic Aneurysm, Abdominal (C0162871), Aneurysm (C0002940), Intracranial Aneurysm (C0007766).

### ATAC-Seq

60,000 cells were collected and washed in PBS. The transposition reaction was performed as described in the Omni-ATAC protocol^66^. Then, the transposition reaction was purified with MinElute PCR Purification Kit (QIAGEN, Hilden, Germany). Tagmented fragments were isolated and PCR-amplified for 6-8 cycles as described previously^36^. Amplified DNA was purified using Agencourt AMPure XP beads (Beckman Coulter, Brea, CA), according to manufacturer’s instructions. ATAC-seq libraries were sequenced using 42 paired-end sequencing cycles on an Illumina NextSeq500 system at the high throughput sequencing core facility of Institute for Integrative Biology of the Cell (CNRS, France, Centre de Recherche de Gif – http://www.i2bc.paris-saclay.fr/). Reads were demultiplexed using bcl2fastq2 (v2.18.12), and 43 to 175 million reads (fragments) were obtained per sample. Adapter sequences were trimmed using Cutadapt (v1.15). Coronary artery and primary SMC and fibroblasts datasets were previously described^16, 36^.

Analyses were performed on the Galaxy webserver^67^. Reads were mapped on GRCh38 (hg38) genome using Bowtie2 v2.3.4.3 with default settings, except reads could be paired at up to 2kb distance. Aligned reads were filtered using BAM filter v0.5.9, keeping only mapped, properly paired reads, and removing secondary alignment and PCR duplicate reads as well as blacklisted regions^68^. ATAC peaks were called using MACS2 callpeak v2.1.1.20160309.6 with default settings. Binary read density files (bigwig) were created using bamCoverage v3.3.0.0.0, normalized on hg38 genome.

To perform sample correlation and principal component analyses, a common list of enriched regions was generated using bedtools multiple intersect (Galaxy Version 2.29.0) in “cluster” mode, and average read coverage on these regions was computed using deepTools multiBamsummary (Galaxy Version 3.3.2.0.0). deepTools plotCoverage and plotPCA functions were used to calculate Spearman correlation between samples and Principal Component Analysis, respectively. Global peak annotation was performed using ChIPSeeker v1.22.0.16. To detect differentially accessible regions between day 0 and day 3, and day 3 and day 8 samples, read counts computed using multiBamsummary were scaled to the total number of reads and used as inputs in DESeq2. Differentially accessible regions were determined using res function and log2 fold changes were determined using lfcShrink function with ashr shrinkage estimator^63^. To compare peaks between day 12 and day 24 (RepSox and TP conditions) or between day 24 (RepSox and TP conditions), primary SMCs and coronary arteries, single peak files were generated using MACS2 callpeak v2.1.1.20160309.6 with pooled inputs from the same condition. Overlaps of peaks were determined using bedtools multiple intersect (Galaxy Version 2.29.0) in “cluster” mode. Motif enrichment analysis was performed using MEME-ChIP tool on MEME webserver (v5.4.1, https://meme-suite.org/), with HOCOMOCO v11 as motif database and default options. Variant enrichment was calculated using rtracklayer (v1.52.1) package as follows. List of lead SNPs were retrieved from GWAS catalog and correspond to individual studies in European populations for coronary artery disease^69^, stroke^70^, blood pressure^71^, intracranial aneurysm^72^, peripheral artery disease^73^, abdominal aortic aneurysm^74^ and migraine^75^. Proxies in high linkage disequilibrium (r^2^>0.7) with the lead SNP in the European population of 1000 Genome reference panel were retrieved at each locus using ldproxy function of LDlinkR package (v1.1.2)^76^. For each peak file, 1000 datasets matched by peak number and sizes for each chromosome were generated. Positions of disease associated variants were converted to hg38 genomic coordinates using UCSC liftover tool (https://genome-store.ucsc.edu/). We counted overlaps of disease associated variants with peak file and background files using countOverlaps function. Variant enrichment was defined as the ratio of the number of overlaps between peak set and average of background sets. Enrichment P-value was evaluated using a binomial test with higher enrichment as alternative hypothesis. P-values were adjusted for the number of tests using Bonferroni method.

### Immunofluorescence staining

Cells were seeded on coverslips in the 24-well plate to approximately 90% confluence. Cells were fixed with 3.7% formaldehyde, permeabilized with 0.5% Triton X-100, and incubated with blocking buffer (PBS supplemented with 0.1% Tween and 5% milk). Cells were incubated with primary antibodies diluted in blocking buffer (1:100). Subsequently, the cells were incubated with Alexa Fluor 488 or 594-conjugated goat anti-mouse or goat anti-rabbit antibodies (1:100) for 1 hour at room temperature and counter-stained with DAPI nuclear dye (1:2500) at room temperature. Coverslips were mounted using mounting buffer and dried for 30 mins at room temperature protected from light. The images were photographed by Zeiss ApoTome.

### Cell viability assay

The cells were seeded at a density of 5,000 cells per well in 96-immuno white immune plates (Thermo Scientific) with 8 duplicates of each cell line. Cell viability was measured by performing CellTiter-Glo 2.0 assay (Promega). Each well was treated with 100 µl of lysis buffer to induce cell lysis and incubated for 30 min at room temperature. The plates were loaded with equal amounts of CellTiter-Glo reagents and the luminescent signal was recorded by using Mithras LB 940 Multimode Microplate Reader machine. Luciferase was measured daily for 5 days starting from the fourth hour after seeding the cells (4h). The viability of all luciferase measurements was normalized to the values at 4 h. Statistical significance was evaluated using Student’s two sample t-test with unequal variances.

### EdU flow cytometry assay

The cells were seeded in 6-well plate at the density of 150,000 cells per well for 1 day. Cells were incubated with EdU (5-ethynyl-2’-deoxyuridine, Life technologies) at final concentration of 10 µM for 4 hours and then harvest to perform Click-iT reaction using Click-iT EdU flow cytometry Alexa Fluor 488 assay kit (Life technologies, Carlsbad, CA, USA, C10420). Fluorescence measurements were taken using LSRfortessa X-20 (BD Biosciences). The results were analysed by Flowjo (BD company). Statistical significance was evaluated using Student’s two sample t-test with unequal variances.

### Transwell assay

The cell invasion assays were performed in a 24-well Transwell chamber with 8µm pore size (Corning). 100 µl of cell suspension (40,000 cells/ml) was added on the top of filter membrane in the Transwell insert and incubated for 10 mins at 37 °C to allow cells to settle down. 600 µl medium with or without chemo-attractant PDGF-BB were added into the lower chamber in a 24-well plate. Plates were incubated for 20 hours at 37 °C. The Transwell inserts were fixed by 70% ethanol and stained with 0.2% crystal violet for 15 min at room temperature. We used ImageJ tool to count and image cells. Statistical significance was evaluated using Student’s two sample t-test with unequal variances.

### Wound Healing Assay

Cells were cultured in 24-well plates to reach confluent monolayers. Straight wounds were performed by 10-ul pipette tips. Plates were washed with media to remove non-adherent cells and photographed for 48 hours with 1-hour intervals at the position of wound gaps. 3 different fields at each time point on each plate were imaged and the wound area was measured by Image J. The wound closure was calculated by the wound area in each period as a percentage of the initial wound area at 0h. Statistical significance was evaluated using Student’s two sample t-test with unequal variances.

### Intracellular Calcium influx measurements

Cells were seeded overnight in 96-well black/clear bottom plates coated with Matrigel (40,000 cells/well). The washing buffer was supplied with 5 mM HEPES and 250 mM probenecide in HBSS with calcium and prepared in the same day. Cells were washed twice and incubated with calcium-sensitive dye Fluo-4-AM (2 µM) (Ex/Em=480 nm/525 nm) for 1 h at 37 °C avoiding exposition to light. Cells were washed twice and incubated in washing solution for 20 mins at room temperature in the dark. The infusion of vasoconstrictor solution including Angiotension ll, PDGF and carbachol was performed by PC-controlled pump. The fluorescence was detected by with 1.52 seconds interval in 180 seconds range. The fluorescence measurements were normalized to the average of first 10 signals.

### Collagen Contraction assay

The cells (2.5×10^5^/well) were harvested and resuspended in culture medium. Collagen gel was prepared by mixing cell suspension and collagen solution and seeded into 45-well plate with Cytoselect Cell contraction Assay Kit (CELL BIOLABS) according to manufacturer’s instructions. The plates were incubated in 37 °C and 5% CO_2_ for 1 hour to allow collagen polymerization. Culture medium (0.5 mL/well) culture medium with or without contraction inhibitor BDM (2,3-Butanedione Monoxime) were carefully added atop each collagen gel lattice. Gels were imaged 20h post-seeding using a RICOH MP3503 scanner (RICOH, Tokyo, Japan). Collagen gel area was determined using Image J software.

### Extracellular matrix collection and protein quantification

Cells were seeded in 10-cm dishes for 48-72 h until 100% confluence was achieved. Cells were then rinsed and incubated in the absence of serum for 1h at 37°C. Cells were washed once by PBS and incubated with 20 mM Ammonium hydroxide at room temperature for 5 mins with gentle agitation to remove all cells. Dishes were washed by copious volume of de-ionized H_2_O four times. The matrix was scraped by 0.1% SDS and frozen in -80°C.

For mass spectrometry analysis, sample were first dried, and proteins were denatured in SDS 2% and TEAB 200mM pH8.5 while disulfide bridges were reduced using TCEP (tris(2-carboxyethyl)phosphine) 10mM and subsequent free thiols groups were protected using chloroacetamide 50mM for 5 min at 95°C. Proteins were trypsin-digested overnight using the suspension trapping S-Trap method (Protifi, Farmingdale, NY, USA) to collect peptides as previously described (Katelyn Ludwig et al., 2018). Eluted peptides were vacuum-dried in a Speed Vac (Eppendorf). LC-MS analyses were performed on a Dionex U3000 RSLC nano-LC system (Thermo Fisher Scientific, Waltham, MA, USA) coupled to a tims-TOF Pro mass spectrometer (Bruker Daltonics GmbH, Bremen, Germany). After drying, digested samples were solubilized in 10 μL 0.1% TFA containing 10% acetonitrile (ACN). One microliter was loaded, concentrated, and washed for 3 min on a C18 reverse-phase precolumn (3 μm particle size, 100 Å pore size, 75 μm inner diameter, 2 cm in length, from Thermo Fisher Scientific, Waltham, MA, USA). Peptides were separated on an Aurora C18 reverse-phase resin (1.6 μm particle size, 100 Å pore size, 75 μm inner diameter, 25 cm in length), connected to a Captive nanoSpray Ionization module (IonOpticks, Middle Camberwell, Australia) with a 60 min run time and a gradient ranging from 99% solvent A, containing 0.1% formic acid in milliQ-grade H2O, to 40% solvent B, containing 80% acetonitrile and 0.085% formic acid in mQH2O. The mass spectrometer acquired data throughout the elution process and operated in the DDA PASEF mode, with a 1.9 s/cycle, with the timed ion mobility spectrometry (TIMS) mode enabled and a data-dependent scheme with full MS scans in PASEF mode. This enabled recurrent loop analysis of a maximum of the 120 most intense nLC-eluting peptides, which were CID fragmented between each full scan every 1.9 s. The ion accumulation and ramp times in the dual TIMS analyzer were set to 166 ms each, and the ion mobility range was set from 1/K0 = 0.6 vs. cm−2 to 1.6 vs. cm−2. Precursor ions for MS/MS analysis were isolated in positive mode with the PASEF mode set to « on » in the 100–1700 m/z range by synchronizing quadrupole switching events with the precursor elution profile from the TIMS device. The cycle duty time was set to 100%, accommodating as many MSMS in the PASEF frame as possible. Singly charged precursor ions were excluded from the TIMS stage by tuning the TIMS using TIMS control software (Bruker Daltonics GmbH, Bremen, Germany).

Identifications (protein hits) and quantifications were performed by comparison of experimental peak lists with a database of theoretical sequences using MaxQuant version 1.6.2.3 (Cox et al. 2014). The databases used were the Human sequences from the NCBInr database (release March 2020) and a list of in-house frequent contaminant sequences. The cleavage specificity was trypsin’s with maximum 2 missed cleavages. Carbamidomethylation of cysteines was set as constant modification, whereas acetylation of the protein N terminus and oxidation of methionines were set as variable modifications. The false discovery rate was kept below 5% on both peptides and proteins.

For differential protein detection, proteins quantified in at least two samples of each condition were kept for further analysis. Label-free quantification intensities from MaxQuant were log2 transformed and normalized using quantile normalization^77^. We performed differential detection analysis using DEqMS package (v1.10.0)^78^. Median peptide counts were used to categorize proteins to compute the variance model.

## Acknowledgements

We would like to thank Pr Ziad Mallat for critical reviewing of the manuscript. This work has benefited from the facilities and expertise of the high throughput sequencing core facility of I2BC (Centre de Recherche de Gif – http://www.i2bc.paris-saclay.fr/) and the 3P5 Proteom’ic platform of Université Paris Cité (https://3p5.recherche.parisdescartes.fr/). The Genotype-Tissue Expression (GTEx) Project was supported by the Common Fund of the Office of the Director of the National Institutes of Health, and by NCI, NHGRI, NHLBI, NIDA, NIMH, and NINDS. Parts of the figure were drawn by using pictures from Servier Medical Art. Servier Medical Art by Servier is licensed under a Creative Commons Attribution 3.0 Unported License (https://creativecommons.org/licenses/by/3.0/).

## Supplemental Materials

Supplementary file 1 (xslx) : RNA-Seq and pathway analysis

Supplementary file 2 (xslx) : ATAC-Seq analysis

Supplementary file 3 (xslx): Underlying data and key ressources used in the study

