## Supplementary Figures for "Genomic, transcriptomic and proteomic depiction of iPSC-derived smooth muscle cells as emerging cellular models for arterial diseases"

* These authors co-supervised this study

#### Supplementary figures legends

**Figure S1. Correlation of gene expression profiles with different cell types**

**A:** Heatmap representation of correlation coefficients between cells isolated at different stages during differentiation (y-axis) and median gene expression profiles in 54 tissues from GTEx database. Pearson correlation coefficient (R) is represented on a white-grey-red scale. Tissues are ranked by the maximum value of R obtained. **B-C:** Correlation of gene expression profiles between cells isolated at different stages during differentiation and gene expression profiles in aorta (**B**) and tibial artery (**C**) samples from GTEx database. Unadjusted Wilcoxon test: ****$p<10^{-4}$.

**Figure S2. Key drivers of differentiation between day 0 and day 3.**

Top differential transcription factor binding motifs in open chromatin regions losing accessibility (downregulated, left panels) or gaining accessibility (upregulated, right panels) between day 0 and day 3 of differentiation. Normalized gene expression of *SOX2*, *NANOG, POU5F1, GATA4, GATA6* and *TEAD1* transcription factors. Binding sites for pluripotent factors *NANOG/POU5F1* and *SOX2* were the most common among regions to lose accessibility during this differentiation process. Consistently, we found an important decrease in the expression for *SOX2* and *NANOG* during the early stage reflecting the loss of pluripotency. Conversely, GATA binding sites were the most common among regions to gain accessibility, with *GATA4* and *GATA6* markedly upregulated, supporting these two transcription factors as potential key regulators at these early stages of SMCs differentiation.

**Figure S3. Key drivers of differentiation between day 3 and day 8.**
Top differential transcription factor binding motifs in open chromatin regions losing accessibility (downregulated, left panels) or gaining accessibility (upregulated, right panels) between day 3 and day 8 of differentiation. Normalized gene expression of *ZIC2*, *ZIC3, JUNB* and *ERG* transcription factors. A substantial reduction in the chromatin accessibility for regulatory elements involving ZIC and GATA binding sites was found at the intermediate stage of differentiation, consistent with the expression profiles of the expression of *ZIC3* and *GATA6* (Supplementary Figure 2). Conversely, binding sites for AP-1 and ETS-like transcription factors gained accessibility, with concomitant increases in the expression of *JUN, JUNB*, *FOS* and ETS-like factor *ERG*, but delayed increases in the expression of *JUN, JUNB*, and *FOS.*

**Figure S4.** Principal component analysis (PCA) of the RNA-seq samples from BW-SMCs, R-SMCs, TP-SMCs and primary SMCs.

**Figure S5.** Violin plot indicating the relative expression distribution of genes associated with gene ontology terms: regulation of VSMC differentiation (A), smooth muscle tissue development (B), SMC migration (C) and SMC proliferation (D) in BW-SMCs, R-SMCs, TP-SMCs and primary SMCs. Unadjusted *P*-value of comparison between sample groups using Wilcoxon test is indicated: *$p<0.05$, **$p<0.01$, **$p<0.001$, ***$p<0.0001$, ns: not significant.

**Figure S6.** Normalized gene expression of extracellular matrix genes *COL1A1*, *COL18A1, COL5A1, FBLN2, FBN1, FBN2, FN1, ELN* and *LOX* in BW-SMCs, R-SMCs, TP-SMCs and primary SMCs.

**Figure S7. A:** Number of proteins quantified from mass spectrometry analysis of each decellularized extract. B: Scatterplot representations of label-free quantification (LFQ) intensity (log2 scale) in TP-SMCs vs R-SMCs. Proteins annotated as extracellular matrix constituents (gene ontology term GO:0031012) are highlighted. Green dots indicate proteins with FDR $< 0.2$ and absolute log2 fold change $> 2$.

**Figure S8.** Distribution in the genome of peaks identified in ATAC-Seq analysis of normal coronary arteries, R-SMCs, TP-SMCs and primary human carotid (HCtASMCs) and coronary (HCASMCs) artery SMCs. Peaks were grouped depending on the groups of samples in which they were identified, as indicated in the left panel.

**Figure S9.** Enrichment (Fold ratio) of disease associated variants in peaks identified in ATAC-Seq analysis of R-SMCs, TP-SMCs, coronary artery SMCs (HCASMC), carotid artery SMCs (HCtASMC) and dermal fibroblasts (HDF). Black outline represents conditions with significant enrichment (*P.adj* $<0.05$).

**Supplementary Figure 1**

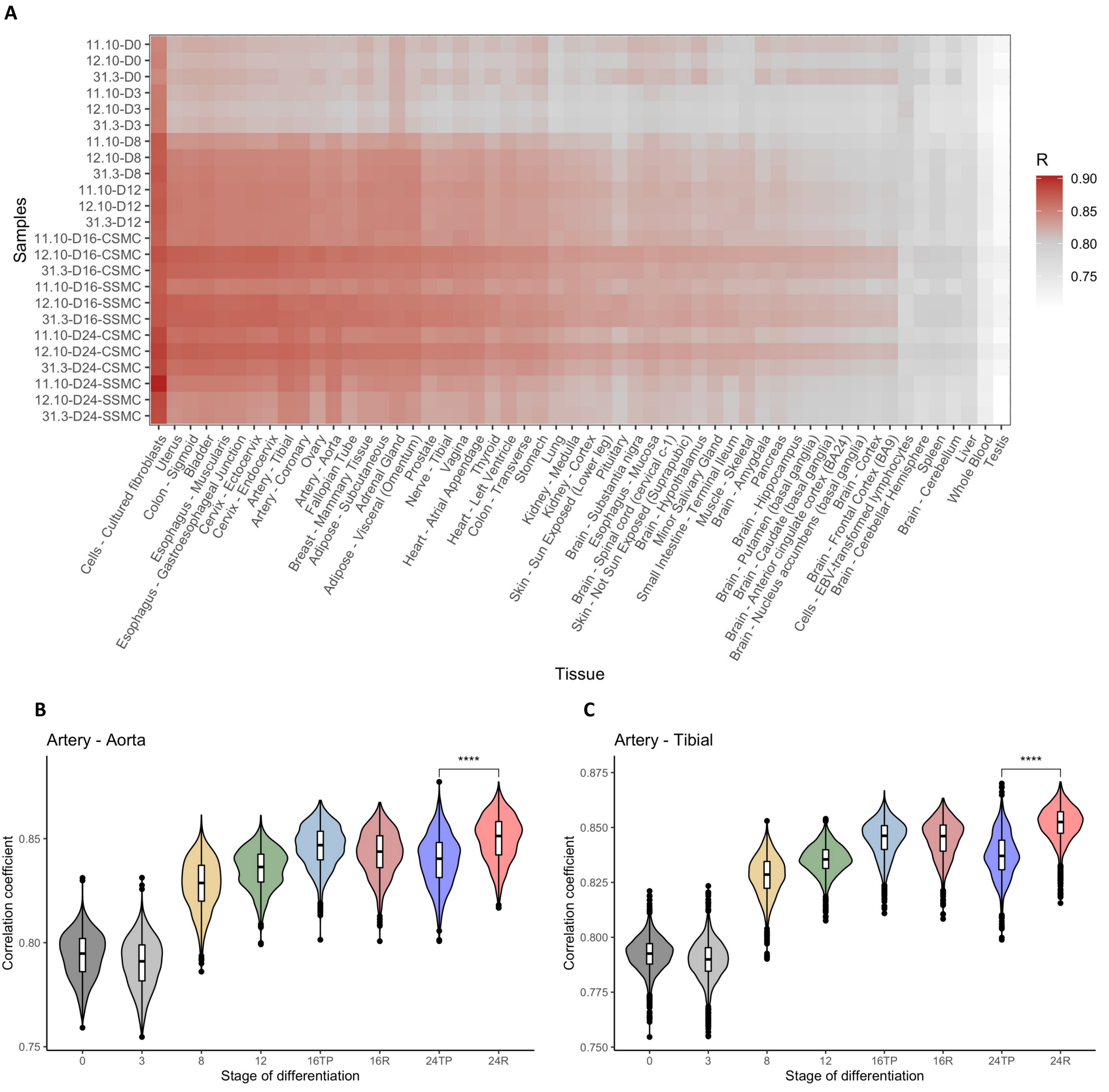

#### Supplementary Figure 2

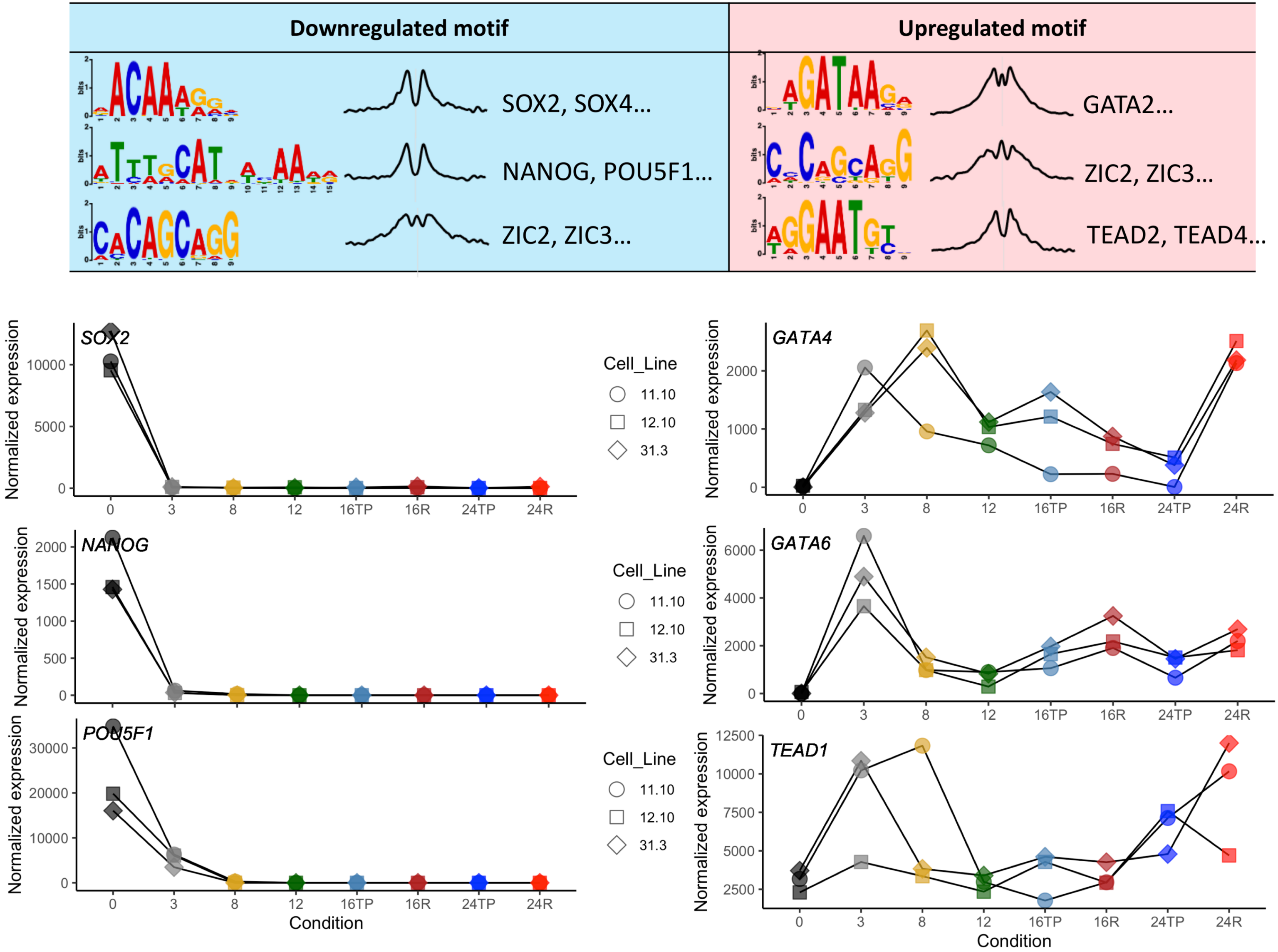

#### Supplementary Figure 3

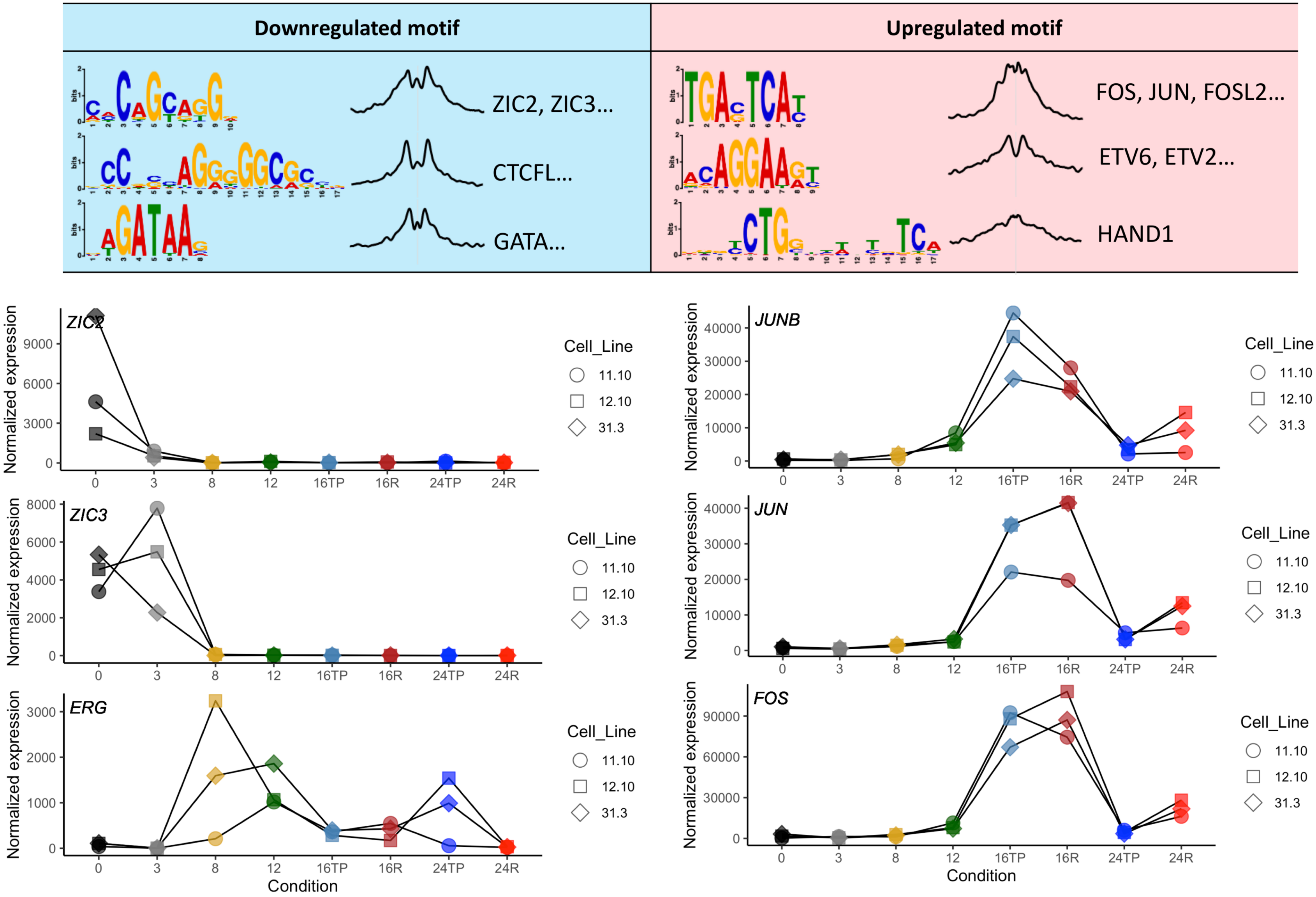

**Supplementary Figure 4**

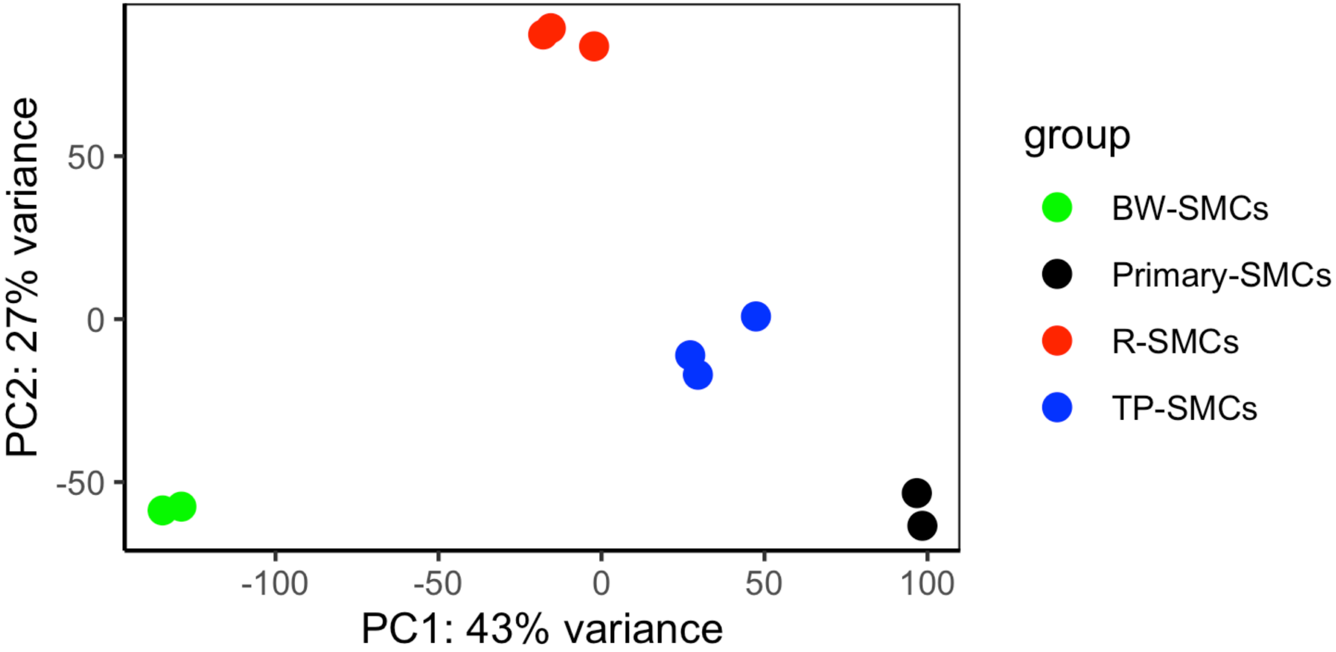

#### Supplementary Figure 5

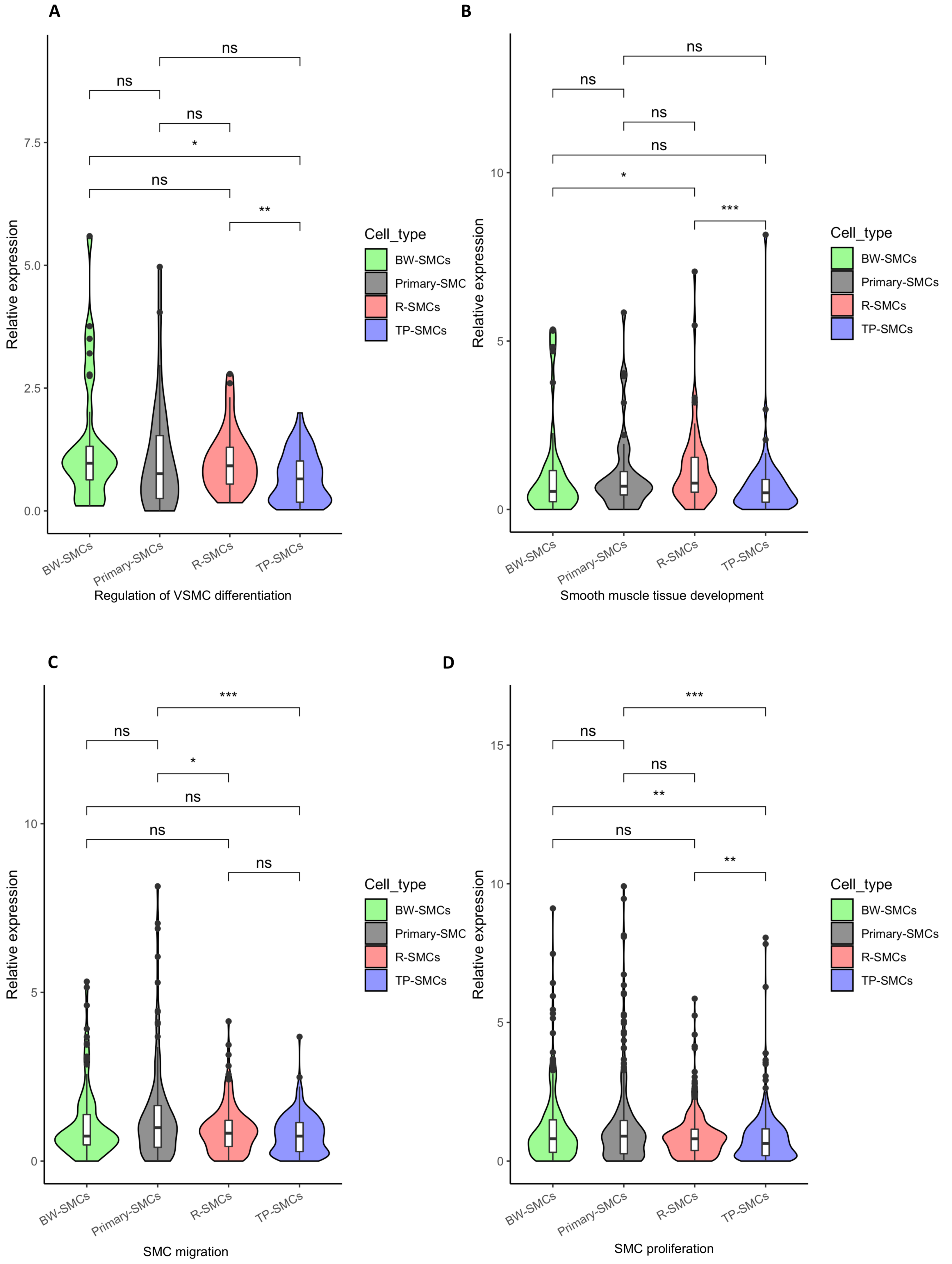

**Supplementary Figure 6**

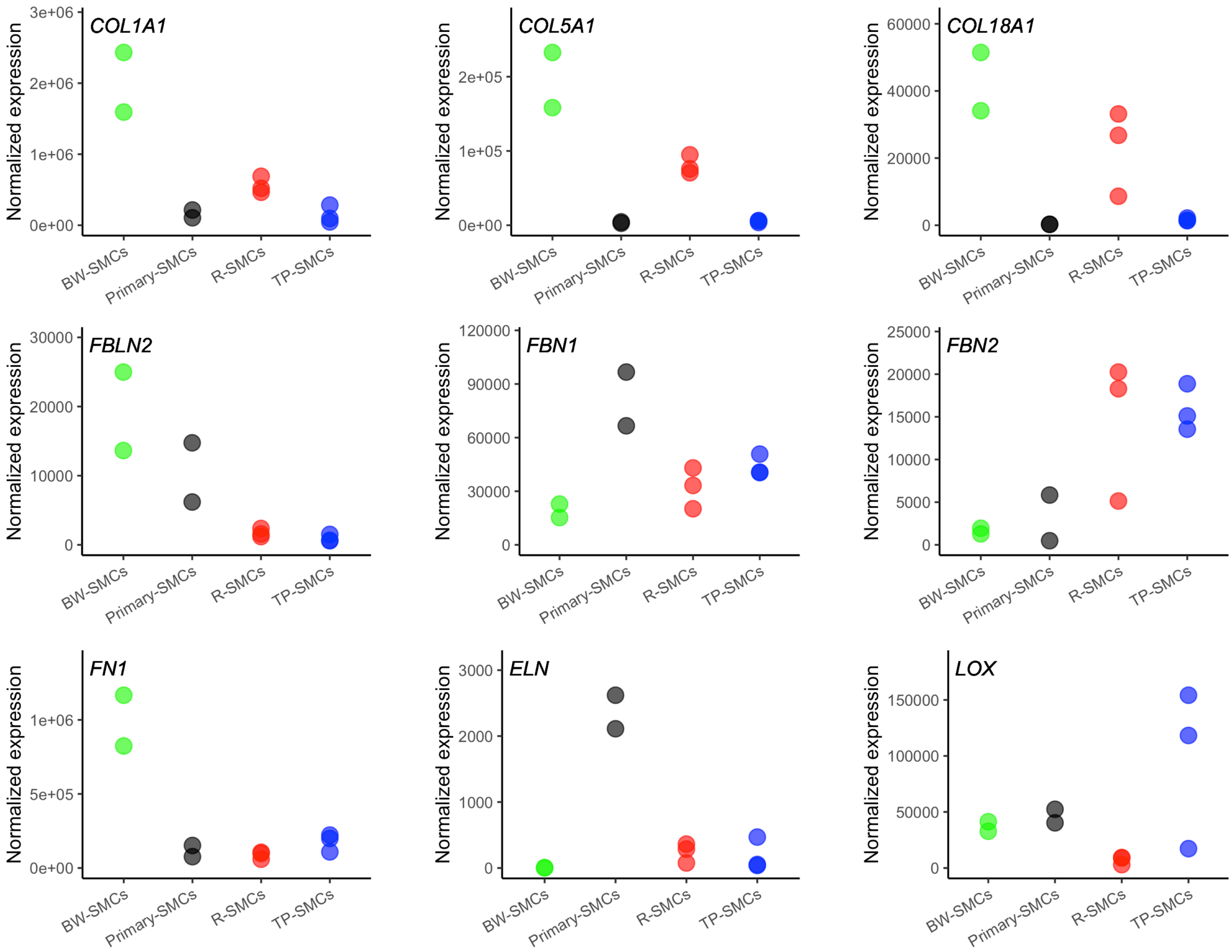

**Supplementary Figure 7**

**A**

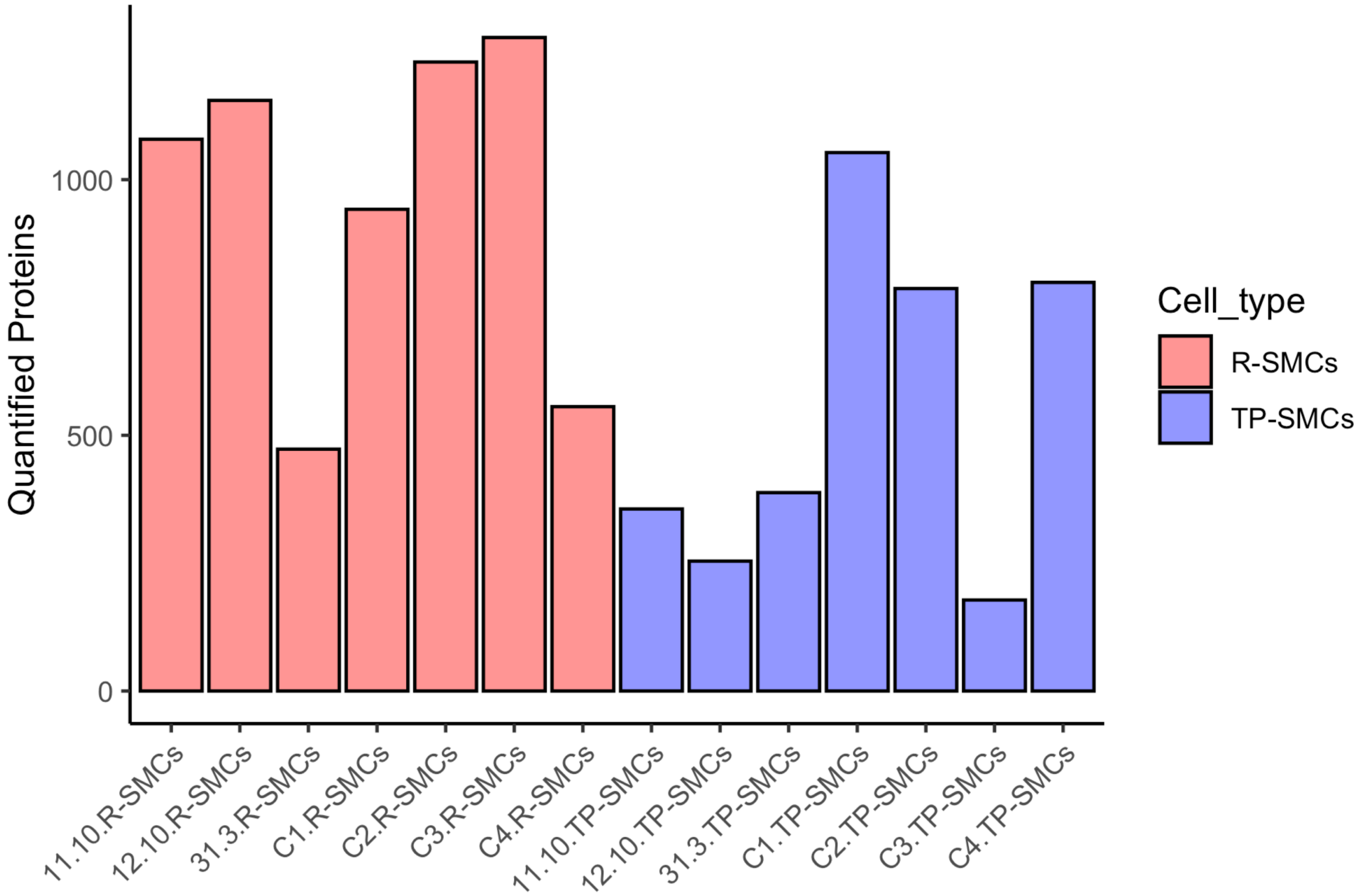


**B**

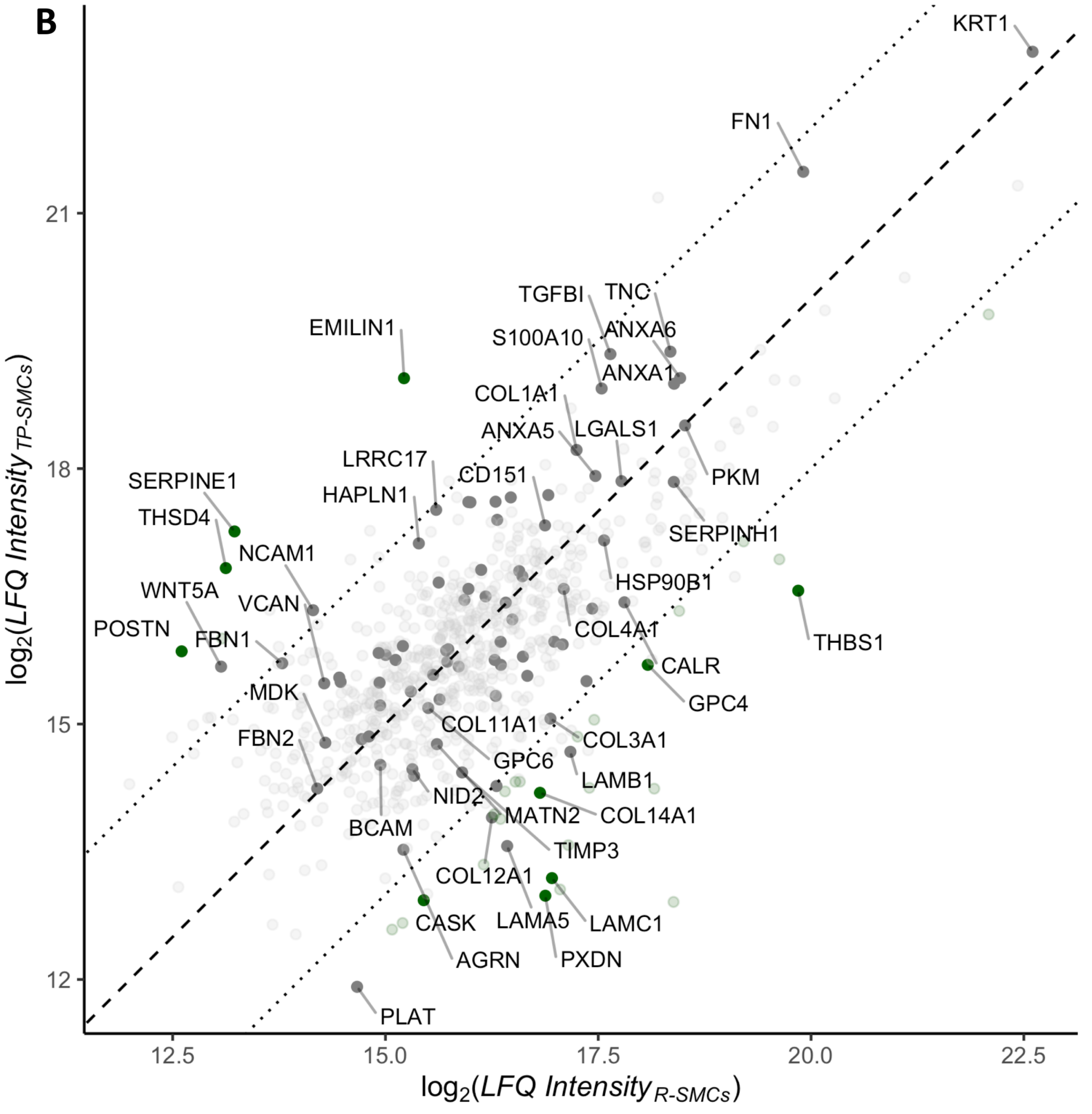

### Supplementary Figure 8

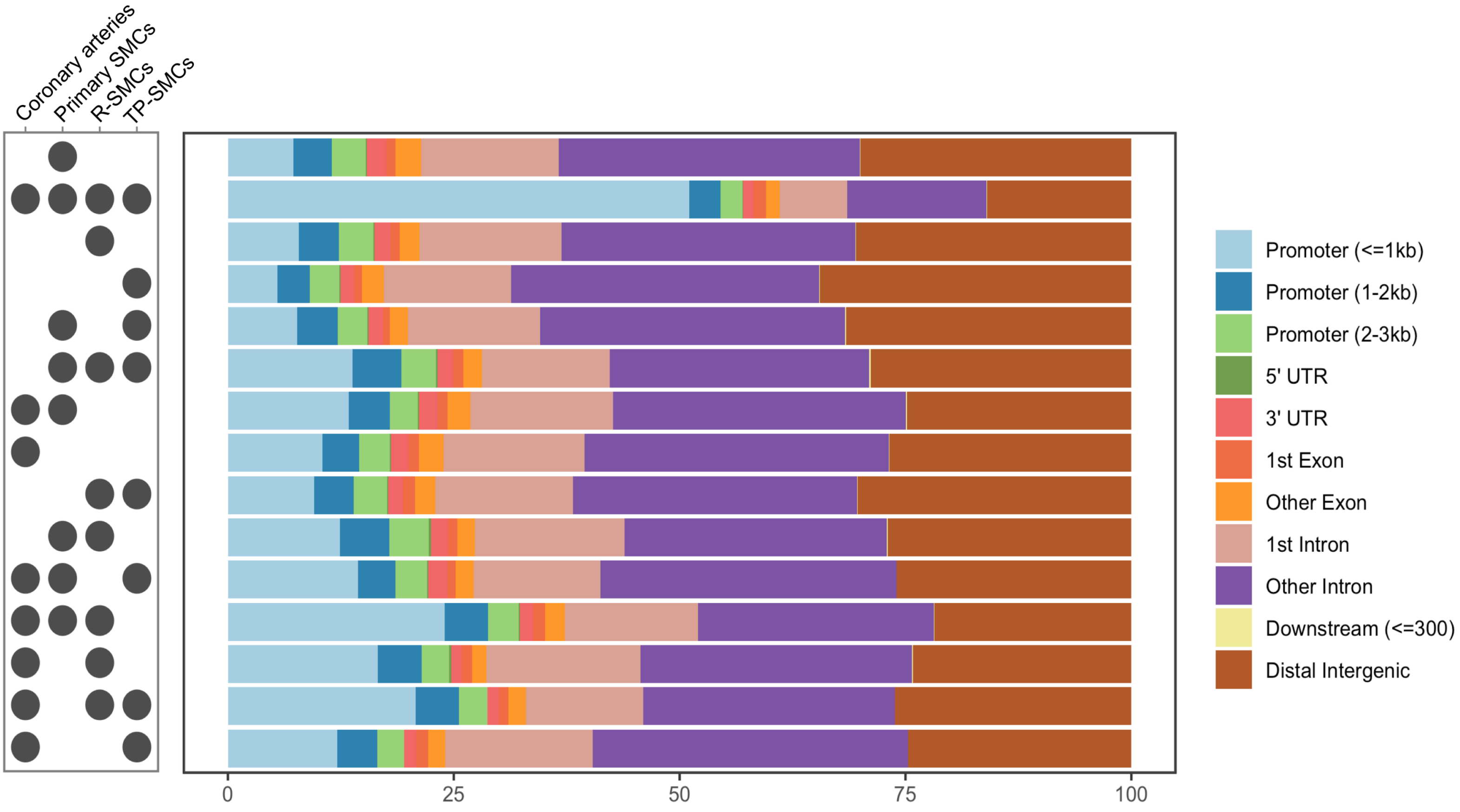

**Supplementary Figure 9**

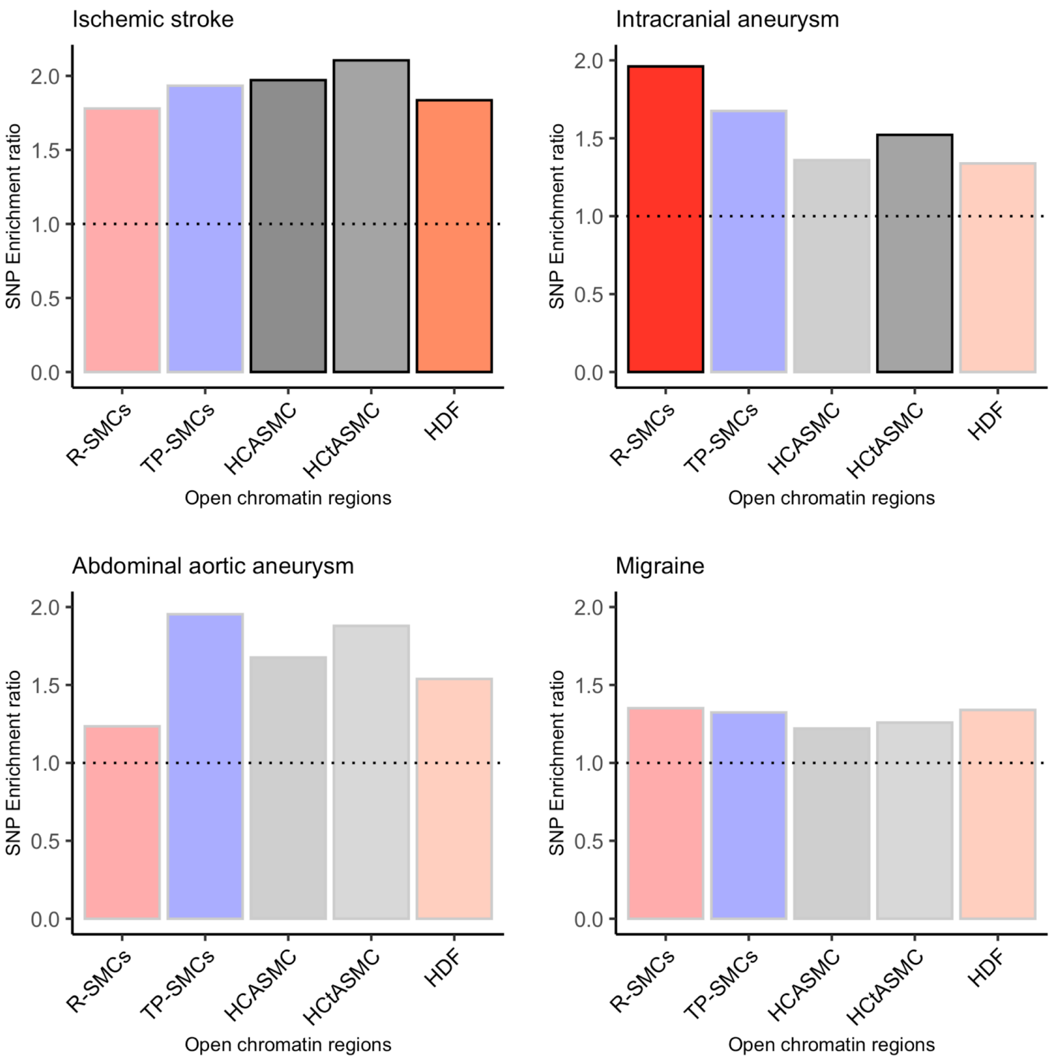
